# A library-based approach allows systematic and rapid evaluation of seed region length and reveals design rules for synthetic bacterial small RNAs

**DOI:** 10.1101/2024.04.24.590872

**Authors:** Michel Brück, Tania S. Köbel, Sophie Dittmar, Adán A. Ramírez Rojas, Jens Georg, Bork A. Berghoff, Daniel Schindler

## Abstract

All organisms must respond to environmental changes. In bacteria, small RNAs (sRNAs) are an important aspect of the regulation network underlying the adaptation to such changes. sRNAs base-pair with their target mRNAs, allowing rapid modulation of the proteome. This post-transcriptional regulation is usually facilitated by RNA chaperones, such as Hfq. sRNAs have a potential as synthetic regulators that can be modulated by rational design. In this study, we use a library-based approach and an oxacillin susceptibility assays to investigate the importance of the seed region length for synthetic sRNAs based on RybB and SgrS scaffolds in *Escherichia coli*. In the presence of Hfq we show that 12 nucleotides are sufficient for regulation. Furthermore, we observe a scaffold-specific Hfq-dependency and processing by RNase E. Our results provide information for design considerations of synthetic sRNAs in basic and applied research.

## Introduction

Bacterial small RNAs (sRNAs) are short regulatory RNA sequences that allow cells to rapidly adapt to environmental changes. sRNAs typically alter the translation of their mRNA targets through imperfect antisense binding, which is mediated by the sRNA seed region. In most cases, sRNAs inhibit translation and decrease the stability of target mRNAs by base-pairing within or in close proximity to the ribosome binding site (RBS) (Holmqvist and Wagner, 2017; Hor et al., 2020; Wagner and Romby, 2015). However, activation of translation by sRNAs is occasionally observed (Frohlich and Vogel, 2009). sRNA-mediated regulation is commonly facilitated by RNA chaperones, with Hfq being one of the best characterized to date. Hfq assembles into a hexameric complex with three RNA interaction sites: the proximal face, the distal face, and the lateral rim (Sauer et al., 2012; Schu et al., 2015; Vogel and Luisi, 2011). sRNAs and mRNAs use distinct RNA sequence motifs to differentially occupy these interaction sites, and Hfq serves as a platform for base-pairing between the RNAs (Santiago-Frangos and Woodson, 2018; Updegrove et al., 2016; Wagner, 2013). With the help of Hfq, the number of nucleotides (nt) that are needed for sRNA-mRNA interactions is often less than ∼10 nt (Santiago-Frangos and Woodson, 2018). In addition, Hfq is important for the stability of sRNAs; Hfq-dependent sRNAs are typically less abundant in the absence of Hfq (Vogel and Luisi, 2011). The majority of mRNA decay in bacteria is initiated by endoribonucleases (predominantly RNase E), and subsequent decay is performed by 5’ and 3’ exoribonucleases that degrade the remnants of RNase E-cleaved mRNA (Bandyra and Luisi, 2018; Hui et al., 2014). The ternary complex of Hfq, sRNA and mRNA may facilitate RNase E-mediated decay of both the mRNA and the sRNA, depending on the 5’ phosphorylation state of the sRNA (Bandyra et al., 2012; Schilder and Gorke, 2023). Hence, RNase E-mediated cleavage considerably contributes to many aspects of sRNA-based regulation (Pfeiffer et al., 2009; Prevost et al., 2011). Because of their regulatory properties, sRNAs are gaining interest as tools to control gene expression and fine-tune phenotypes of bacterial cells.

Synthetic sRNAs have tremendous potential to expand the toolbox of genetic engineering techniques, allowing for strain improvement to increase the yield of the target product (Na et al., 2013). Synthetic sRNAs have the advantage of allowing rapid plasmid-based assays (Köbel et al., 2022) to identify efficient target regulation prior to performing labor-intensive genome engineering procedures. While synthetic sRNAs are promising tools for genetic engineering, antisense peptide nucleic acids (PNAs) have the potential to serve as on-demand programmable antibiotics by targeting essential or resistance genes of specific target organisms (Oh et al., 2014; Popella et al., 2022; Popella et al., 2021). Unlike an antibiotic, such PNAs can be rapidly modified by changing the antisense sequence, which has enormous potential to address the antibiotic resistance crisis. Furthermore, species-specific targeting might be achievable with such an antisense strategy, leaving the majority of the microbiome intact (Vogel, 2020). For these reasons, computational tools have been developed to predict the optimal seed region for antisense binding of synthetic sRNAs or antisense oligomers (Jung et al., 2023; Philipp et al., 2023). Such prediction tools allow systematic testing but do not guarantee sRNA functionality (Brück et al., 2024). The binding probability can be estimated by calculating the binding energy, an approach that depends on careful consideration of structural RNA features (Mann et al., 2017). It is generalized that the lower the binding energy the stronger the binding between two RNA species. However, in addition to the binding energy, other factors such as the accessibility and location of the antisense-mediated binding region and the secondary structures of the synthetic sRNA or mRNA may play important roles. There is a lack of knowledge to predict and apply perfect synthetic sRNAs on demand. Systematic studies are needed to improve prediction tools and to generate training data for artificial intelligence-based prediction in the future for time and cost effective application of synthetic sRNAs in basic and applied research.

Here, we use a library-based Golden Gate cloning approach and laboratory automation workflow to obtain a library of synthetic sRNAs with increasing seed region lengths (SRL) and to characterize the functionality of these synthetic sRNAs in *Escherichia coli*. The experimental design is based on our previously characterized synthetic sRNAs and an established oxacillin susceptibility assay (Köbel et al., 2022). We use the designed library to identify the minimum SRL for functional synthetic sRNAs. We investigate the steady-state levels of synthetic sRNAs, the potential for Hfq-independent post-transcriptional regulation with increasing SRL and RNase-dependent processing of selected candidates. We further extend the library-based approach from the RybB sRNA containing a 5’ seed region to the structurally complex SgrS sRNA containing a 5’ open reading frame (*sgrT*) and an internal seed region. Our results show that the SRL differentially affects RybB and SgrS. RybB functionality seems to be independent of the SRL, albeit RNase E-mediated processing occurs with increasing SRL. In case of SgrS, such processing is not observed. Instead, SgrS functionality is sensitive to SRL-mediated changes in local secondary structures, and an increasing SRL helps to alleviate the Hfq-dependency of SgrS. These observations underscore that library-based approaches provide efficient means to optimize synthetic sRNA design and to study sRNA biology.

## Results

### Concept of Golden Gate-based sRNA library construction and validation

sRNAs have a modular structure and can be as simple as a 5’ seed region, capable of binding complementary mRNAs, and a short single-stranded region followed by a stem-loop terminator structure (Figure 1A). Their small size and the modular structure allow the standardized, high-throughput construction of synthetic sRNAs to be tested and used as post-transcriptional regulators in synthetic biology and biotechnology applications. We have previously described an easy-to-use plasmid toolset for efficient generation and benchmarking of synthetic small RNAs in bacteria (Köbel et al., 2022). This has been the basis for the development of a Modular Cloning toolbox utilizing SapI, an enzyme generating 3-bp overhangs and thereby reducing the size of the fusion sites compared to the 4-bp overhangs that are generated by other Type IIS restriction enzymes (e.g., BbsI, BsmBI, BsaI) (de Vries et al., 2024; Weber et al., 2011). Briefly, this system allows the generation of level 0 constructs which can be combined to transcription units (TU) in designated acceptor plasmids with minimal fusion sites reducing potential scar sequences (Figure 1B). For this study we created a pBAD-derived acceptor plasmid for cloning of distinct sRNA TUs via SapI (Brück et al., 2024). The plasmid has the same properties and features as in our previous study, including the *mCherry-ccdB* counter-selection cassette, only the Type IIS enzyme was swapped from BbsI to SapI (Köbel et al., 2022). In our previous study, we tested a set of 16-nt seed regions in combination with the RybB scaffold to repress *acrA* mRNA which allows us to perform oxacillin susceptibility assays to validate sRNA functionality (*acrA* repression leads to an increase in oxacillin susceptibility). We picked 16-nt seed regions because it resembles the length of the natural RybB seed region (Papenfort et al., 2010). However, for synthetic biology applications we were wondering about the effects of SRL variation and its correlation to post-transcriptional regulation. We created a library based on the previously described 16-nt seed region s8 (s8 = seed sequence 8 of the initial screen; *cf*. (Köbel et al., 2022)), with seed regions ranging from 2 to 82 nt in length and 2-nt increments towards the transcription start site (TSS) of *acrA* mRNA (Figures 1C and S1). The corresponding sRNAs were denoted SRL2 to SRL82, respectively. The 5’ untranslated region (UTR) of *acrA* was previously determined to have a length of 79 nt (Eguchi et al., 2003). Notably, we were expecting that based on the cloning fidelity the smaller seed regions will not be generated. The SRL-library was ordered as an oligonucleotide pool, in which each individual SRL-oligonucleotide contained overhangs for PCR amplification and SapI recognition sites for Golden Gate cloning (Figure 1C). To create the plasmid library, the level 0 parts for the promoter, the scaffold, the destination plasmid, and the PCR-amplified library are combined in one Golden Gate reaction. The reaction is directly transformed into *E. coli* MG1655 wild-type cells.

**Figure 1.**
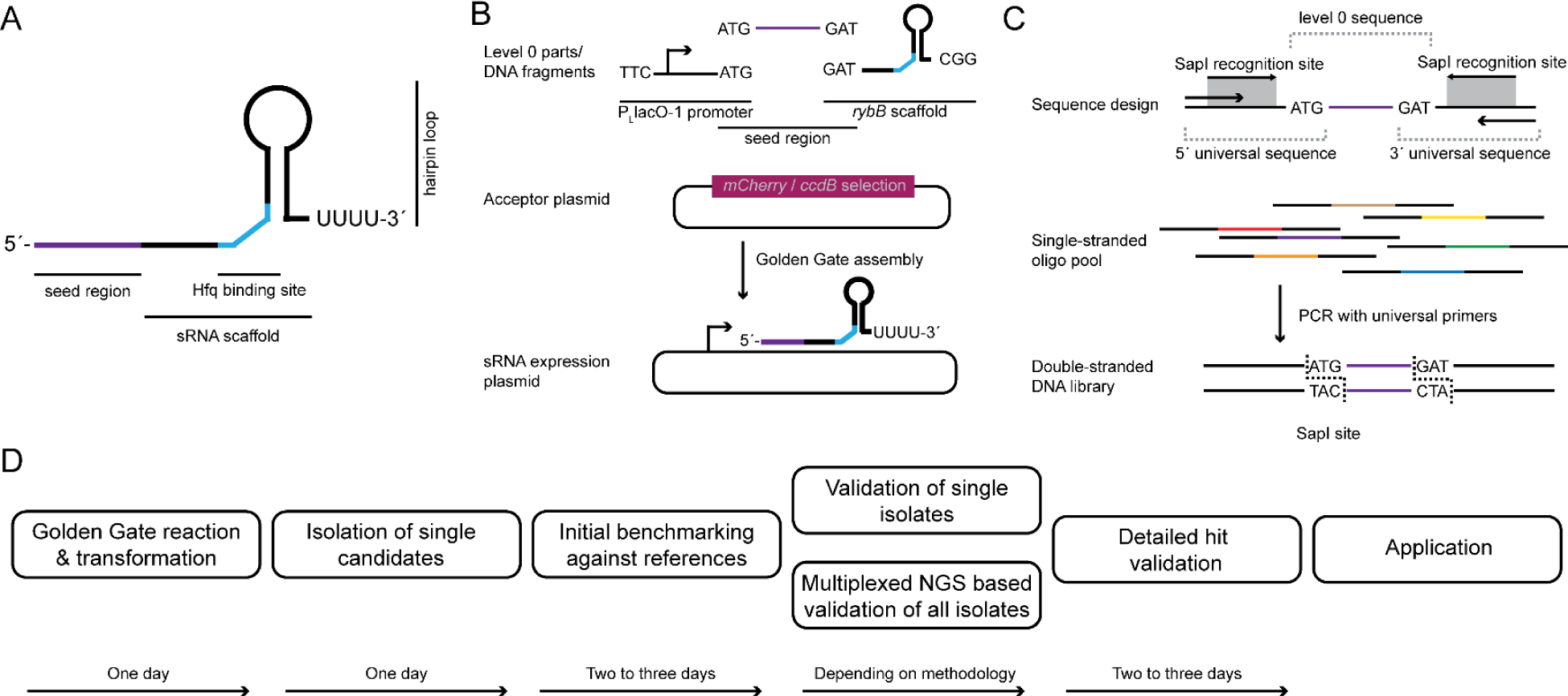
Workflow of synthetic sRNA cloning, library generation and characterization. **(A)** Synthetic sRNAs for post-transcriptional control can have a short length and a simple structure consisting of the 5’ seed region and the sRNA scaffold. The seed region can be modified for specific targeting of mRNAs. **(B)** Based on the simple structure, sRNAs can be modularized and constructed by Golden Gate cloning. The transcription unit of the constructed synthetic sRNAs consists of the promoter (in this case PLlacO-1), a seed region and the sRNA scaffold (in this case RybB). The three parts are combined into the appropriate acceptor plasmid for subsequent experiments. **(C)** Concept of oligonucleotide pool-based libraries. The pooled oligonucleotide sequences contain 5’ and 3’ universal sequences providing the required Type IIS recognition and cut sites (the Type IIS recognition site is visualized by the gray box with arrows for its direction and the cut site is highlighted by the indicated nucleotide sequence of 3 nts in length). The universal sequences serve as primer binding sites (indicated by the outermost arrows) to convert the single-stranded oligonucleotide pool into a double-stranded DNA library that can subsequently be used for cloning approaches. Different colors of oligo pool sequences indicate different seed region lengths in the library. **(D)** Schematic workflow for construction and selection of synthetic sRNAs. The workflow can be completed within < 10 days depending on the chosen sequence validation method to obtain characterized synthetic sRNAs for their application.

For subsequent characterization, we have established a workflow supported by automated equipment to increase throughput (Figure 1D). This workflow is designed to isolate individual candidates from the library and perform an initial screening. Working with a pooled library is not possible in this case because functional candidates have reduced viability under our chosen selection pressure, the β-lactam antibiotic oxacillin. We also tested whether a rapid workflow for colony PCR-based sequencing could be established using Nanopore sequencing to identify individual candidates (Ramirez Rojas et al., 2024a; Ramirez Rojas et al., 2024b). To achieve this in a cost-effective and sustainable manner, we used our previously described highly multiplexed barcoding approach and downscaled reaction volumes using a combination of a bulk dispenser, an acoustic dispenser, and a screening robot to minimize reagent use and plastic waste. Using this workflow, 66% of the candidates could be assigned a SRL. For the remaining 34%, either coverage was insufficient, the result was ambiguous (possibly not a single isolate), or no amplicon was obtained. Using this protocol, four experiments were sequenced simultaneously on one Flongle cell, resulting in 1,536 tested candidates, resulting in a cost of less than 0.10 € per barcoded amplicon, including colony PCR, sequencing library generation and Nanopore sequencing (Ramirez Rojas et al., 2024a).

### Functional characterization of the synthetic SRL-library using the RybB scaffold

To create a RybB library with varying SRL, two level 0 plasmids containing either the P_L_lacO-1 promoter (hereafter, PL) (pSL135) (Pfeiffer et al., 2009; Vogel et al., 2004) or the RybB scaffold sequence (pSL123), the destination plasmid (pSL137), and the amplified SRL-library were assembled in one Golden Gate reaction. The reaction was transformed into *E. coli* wild type MG1655 (hereafter referred to as “wild type”), and dilutions were plated to obtain plates with optimal density for automated colony picking. For the purpose of workflow establishing, 380 candidates were picked into a 384-well format (Figure 2A). The last four positions were kept empty to accommodate controls. The processing of the samples was performed according to the described workflow (Figure 1D). In a first step, the 380 candidates and four controls were screened on solid media containing varying concentrations of oxacillin (0 to 200 µg/mL) (Figure 2A, not all data shown). The controls are the wild type with an empty plasmid, the Δ*acrA* strain with an empty plasmid, the previously characterized pP_L_-RybB-s8 strain (s8 = 16-nt seed region from (Köbel et al., 2022)), and a media control. Strains were cultivated in the 384-well format overnight and spotted onto solid media for the initial screen (Figure 2A). Under the reference condition, without oxacillin, all patches show growth. Under the testing condition with 75 µg/mL oxacillin, however, growth is reduced for several candidates. Importantly, the previously characterized pPL-RybB-s8 control is showing no strong phenotype, indicating that the SRL-library contains candidate sRNAs with a superior post-transcriptional regulation. Based on these results, candidates which show increased oxacillin susceptibility could be individually analyzed by Sanger sequencing. However, we were interested in the composition of the whole library. Therefore, we decided to sequence all candidates in house on the Nanopore sequencing platform with a custom dual barcoded amplicon sequencing procedure (*cf*. Methods) (Ramirez Rojas et al., 2024a; Ramirez Rojas et al., 2024b). Based on the results, we successfully generated an SRL-collection from 2 to 82 nt incrementing in 2-nt steps. In addition, the library contained a control with no seed region at all (denoted “SRL0”). Three constructs with an SRL of 4, 8, and 12 nt were not obtained within the library, presumably due to low cloning fidelity of short fragments. We used the defined SRL-library to repeat the screening on solid agar by spotting dilution series onto medium with increasing oxacillin concentration for a qualitative susceptibility assessment (data not shown). For quantitative evaluation we performed plate reader growth kinetics in five replicates to determine the area under the curve (AUC) for susceptibility assessment (Figure 2B). It is not surprising that with an increased SRL the binding energy decreases, as assessed by IntaRNA predictions (Mann et al., 2017). However, we did not observe a linear increase of post-transcriptional regulation for the synthetic sRNAs with increasing SRL (Figure 2B). Furthermore, regulation strength was not correlated with the expression level of sRNAs, as indicated by Northern blot analysis (Figure S2).

**Figure 2.**
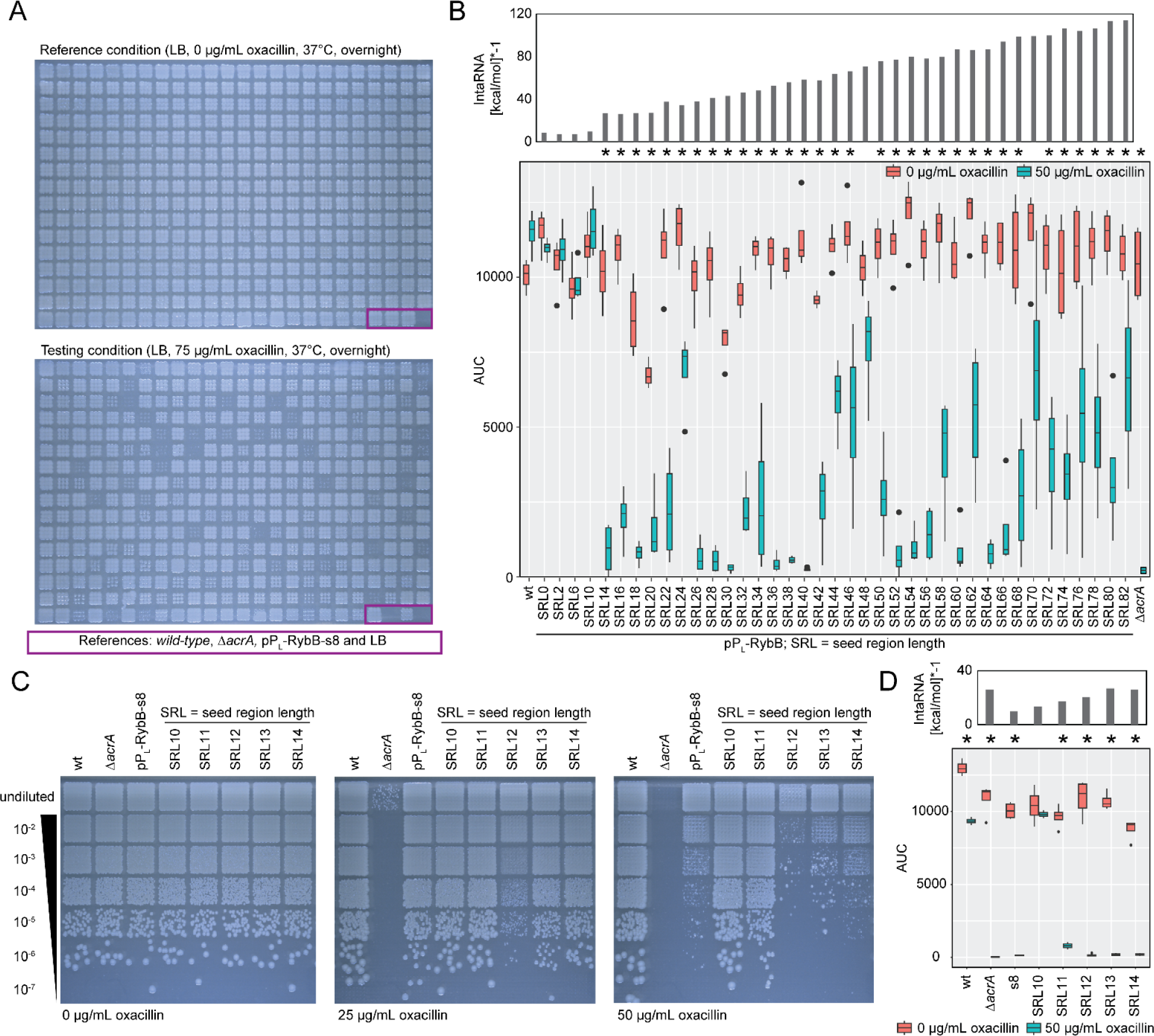
Initial screening and detailed characterization of the RybB SRL-library targeting *acrA* mRNA. **(A)** Screening of RybB SRL-library. 380 candidates and four respective controls were spotted from fresh overnight cultures into 3 × 3 grits on LB-agar plates with and without oxacillin. Comparing the reference and the testing condition indicates multiple candidates render cells susceptible towards oxacillin. Comparing the established pPL-RybB-s8 construct (Köbel et al., 2022) with the library indicates sRNA constructs with a superior regulation. Violet box indicates the position of indicated controls. **(B)** Liquid growth analysis of the SRL-library indicates regulation for the majority of synthetic sRNAs by comparing the area under the curve (AUC) in the presence and absence of oxacillin. According to the assay, at least 14 nt are needed for regulation of *acrA* mRNA. Further, the SRL does not correlate with its functionality despite possessing lower binding energies. Binding energies were calculated by IntaRNA (for display reasons the values are multiplied by -1). Wild type (wt) and Δ*acrA* with an empty plasmid serve as positive and negative control, respectively. Oxacillin susceptibility assay was performed in quadruplicate. Student’s *t*-test was applied for statistical testing of the difference between growth with and without oxacillin (*: *P* < 0.05). **(C)** Oxacillin susceptibility test on solid medium reveals that 12 nt seem to be the minimal SRL for functionality of synthetic sRNAs. A representative replicate is shown. **(D)** Oxacillin susceptibility assay in liquid culture shows similar results. However, in liquid media a SRL of 11 nt already shows regulation, presumably based on the increased forces onto the bacterial cell wall. Binding energies were calculated by IntaRNA (for display reasons the values are multiplied by -1). Oxacillin susceptibility assay was performed in quadruplicate. Student’s *t*-test was applied for statistical testing of the difference between growth with and without oxacillin (*: *P* < 0.05). pPL-RybB-s8 is abbreviated as s8.

Based on our obtained results, we identified an SRL of 10 nt to not be functional in our assay, while an SRL of 14 nt was functional (Figure 2C-D; note: s8 = 16-nt seed region as control from (Köbel et al., 2022)). To dissect the threshold for our assay and determine if there are quantitative differences, we individually generated sRNA constructs with an SRL of 11, 12 and 13 nt, and performed their characterization on solid and in liquid media in comparison to the respective controls. An SRL of 11 nt showed a slight post-transcriptional regulation which significantly improved with a length of 12 nt (Figures 2C-D). It is, however, possible that these observations are specific for the targeted mRNA sequence and may differ in regard to other targets. The results indicate a stronger regulation in liquid culture which is most likely caused by increased forces onto the bacterial cell wall making oxacillin under this condition more potent.

We selected the synthetic RybB sRNAs with an SRL of 12, 16, 30, 40, 52, and 64 nt for further analyses. The sRNAs were selected because they represent (i) the minimal length needed for regulation (SRL12), (ii) the reference length from our previous study (SRL16), (iii) the best performing (SRL30 and 40), or (iv) well-performing sRNAs with an SRL of > 50 nt (SRL52 and 64).

### Synthetic RybB sRNAs rely on Hfq irrespective of the seed region length

It was previously reported that RybB is associated with Hfq (Sauer and Weichenrieder, 2011; Thompson et al., 2007). Having the comprehensive SRL-library on hand, we were wondering if increasing the SRL will reduce Hfq-dependency of synthetic RybB sRNAs. The defined SRL-library was transformed into an Δ*hfq* strain and subsequently analyzed on solid agar (data not shown) and in liquid culture (Figure S3). We observed that the Δ*hfq* strain shows generally a higher susceptibility towards oxacillin which was confirmed by testing the minimum inhibitory concentration (MIC); the MIC of the Δ*hfq* strain was reduced approximately 4-fold in contrast to the wild type (Figure S4). The oxacillin concentration was, therefore, adjusted for experiments with the Δ*hfq* strain. For our detailed analysis we focused on the subset of functional RybB sRNAs chosen in the previous section (SRL of 12, 16, 30, 40, 52, and 64 nt). The comparative growth on solid and in liquid media showed the expected oxacillin-sensitive phenotype in the wild type but not in the Δ*hfq* strain (Figure 3A, Figure S5). To validate the results on the level of post-transcriptional regulation, we tested the selected sRNAs using an established fluorescent reporter construct; a translational fusion of the first nine amino acids of *acrA* with *syfp2* (Köbel et al., 2022). As expected, all selected sRNAs caused an at least 10-fold reduction of sYFP2 fluorescence in the wild type. In the Δ*hfq* strain, the fluorescence was only reduced by approximately 2-fold, irrespective of the SRL (Figure 3B). We conclude that (i) the SRL does not influence regulation strength when using the RybB scaffold and (ii) that all tested synthetic RybB sRNAs are Hfq-dependent. To further explore the Hfq-dependency, expression levels of the selected sRNAs were analyzed on Northern blots, which revealed the expected sizes of the sRNA primary transcripts in both the wild type and the Δ*hfq* strain (Figure 3C). However, steady-state levels in the Δ*hfq* strain were decreased, indicating a reduced stability of the synthetic RybB sRNAs, which supports the role of Hfq as a major determinant for sRNA stability (Vogel and Luisi, 2011). Interestingly, Northern blot experiments revealed distinct processing patterns, which were especially apparent for longer seed regions (Figure 3C). Notably, the processing pattern seems not to show a positive or negative influence towards sRNA functionality.

**Figure 3.**
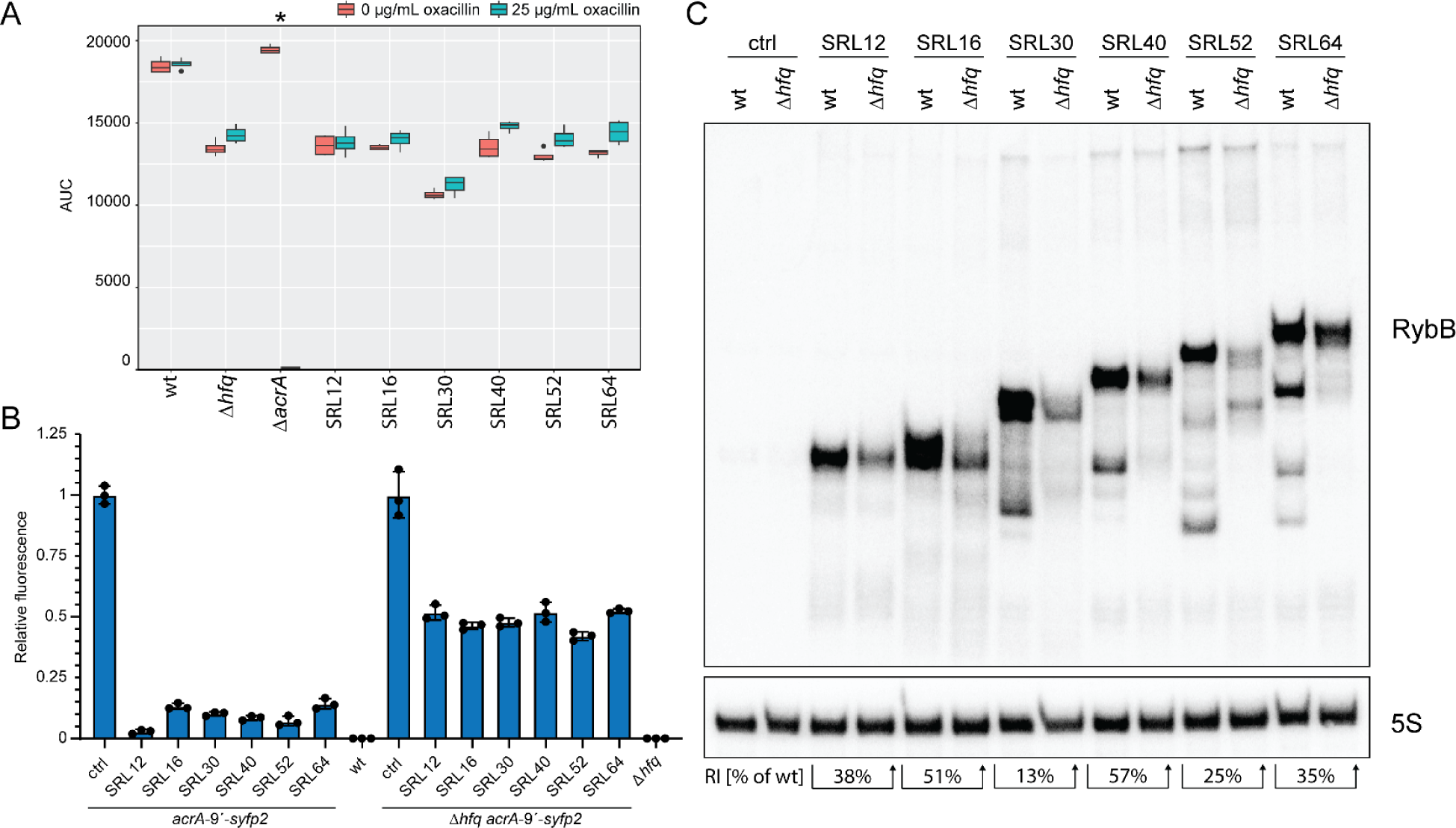
Hfq is required for functionality of synthetic RybB sRNAs. **(A)** Liquid growth oxacillin susceptibility assay for selected RybB candidates in Δ*hfq*. No regulation of the *acrA* target can be observed, which is consistent with the solid media susceptibility assay (Figure S5). The analysis indicates a reduced viability of Δ*hfq* in contrast to the wild type (wt) and Δ*acrA*, already in the absence of oxacillin. Oxacillin susceptibility assay was performed in quadruplicate. Student’s *t*-test was applied for statistical testing of the difference between growth with and without oxacillin (*: *P* < 0.05). **(B)** The functionality of selected RybB candidates is validated by a previously established sYFP2-reporter assay using the chromosomal *acrA-9’-syfp2* construct (Köbel et al., 2022). The sRNAs show functionality in the presence of Hfq. In the absence of Hfq a fluorescence reduction of 50% can be observed for all sRNAs irrespective of the SRL. The control (ctrl) refers to empty plasmid pSL009. A wild type (wt) and Δ*hfq* strain without reporter construct were used as background controls, respectively. Fluorescence reporter assay was performed in triplicate. **(C)** Comparison of abundance and processing of selected RybB sRNAs in wild type (wt) and Δ*hfq* by Northern blot analysis. The control (ctrl) refers to empty plasmid pSL009. 5S rRNA serves as a loading control. Abundance of RybB sRNAs is reduced on average to 37% +/- 16% (RI = relative intensity in Δ*hfq* in comparison to wild type).

### RybB sRNAs from the SRL-library are predominantly processed by RNase E

Two major factors for sRNA processing are the endoribonucleases RNase III and E. RNase III cleaves double-stranded RNAs and is required for the processing of non-coding RNAs (Bechhofer and Deutscher, 2019; Court et al., 2013; Robertson and Dunn, 1975). We, therefore, tested the selected synthetic RybB sRNAs in an RNase III deletion background and its isogenic wild-type strain. On Northern blots, the processing pattern of the synthetic sRNAs was similar in both strains (Figure 4A). However, we observed a slight increase of sRNA abundance (both primary transcript and processing products) in the RNase III deletion background, suggesting a reduced sRNA decay in this strain, which is in line with the general function of RNase III (Bechhofer and Deutscher, 2019; Viegas et al., 2007). RNase E is an essential RNase which cleaves single-stranded RNAs and plays a prominent role in sRNA processing and decay (Chao et al., 2017; Masse et al., 2003; Quendera et al., 2020). We investigated the processing pattern of the selected sRNAs in a temperature-sensitive RNase E background under permissive (30°C) and non-permissive (44°C) conditions. Our results indicate that RNase E is at least one of the RNases responsible for the observed processing (Figure 4B). The processing pattern is visible under permissive but much less pronounced under non-permissive conditions. When inspecting the seed sequences, we observed potential RNase E cleavage sites as AU-rich motives (Chao et al., 2017). The potential processing sites correspond with our experimental data (Figure 4C and S2). Processing of the synthetic RybB sRNAs may occur only after binding to its target mRNA. We, therefore, investigated potential alterations of the processing pattern in a Δ*acrA* strain. However, we did not observe any differences in comparison to the wild type and conclude that processing is at least independent of *acrA* (data not shown). In summary, we conclude that the synthetic RybB sRNAs are predominantly processed by RNase E, probably due to AU-rich motives in the seed regions. This finding may have important implications for synthetic sRNA design, in particular if target mRNAs have a high AU-content in the target region.

**Figure 4.**
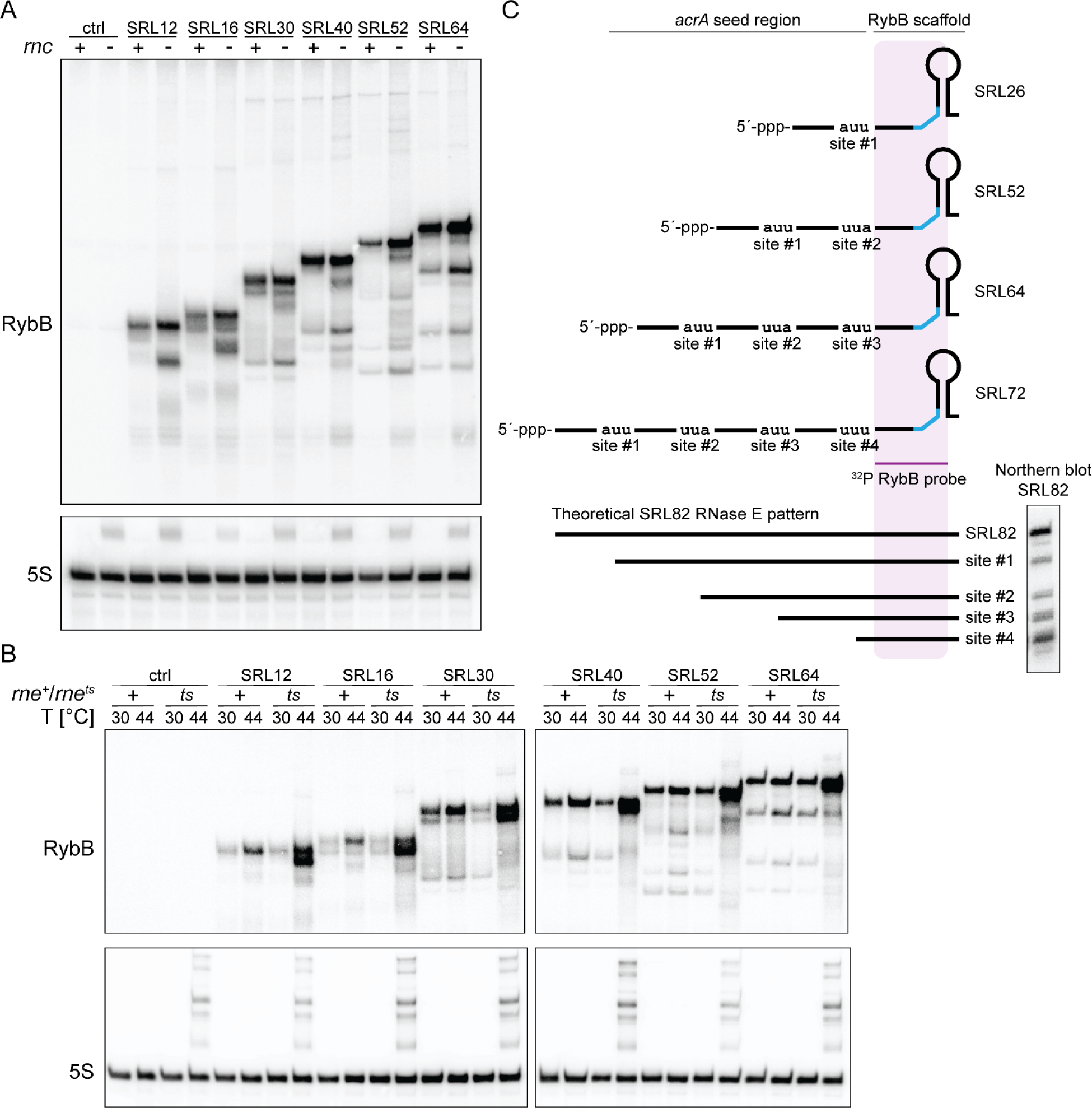
Investigation of RNase-mediated processing of synthetic RybB sRNAs. **(A)** Northern blot analysis of synthetic RybB sRNAs in an RNase III deletion strain (*rnc*^-^) and an isogenic wild type (*rnc*^+^). The control (ctrl) refers to empty plasmid pSL009. 5S rRNA serves as a loading control. **(B)** Northern blot analysis of synthetic RybB sRNAs in a temperature-sensitive RNase E background (*rne^ts^*) and an isogenic wild type (*rne*^+^). The control (ctrl) refers to empty plasmid pSL009. 5S rRNA serves as a loading control. **(C)** Distinct processing patterns with stable intermediates can be observed which matches potential RNase E cleavage sites (Chao et al., 2017). SRL26, SRL52, SRL64, and SRL72 are visualized because of the first appearance of the processing band in the Northern blot of the whole SRL library (see Figure S2). Theoretical and detected processing pattern exemplary visualized for RybB SRL82. For the full Northern blot panel of RybB SRL library see Figure S2.

### Applying the SRL-library approach to the more complex SgrS sRNA

RybB has a rather simple structure. However, there are sRNAs which have more complex structures, as for example SgrS which has a length of 227 nt. Based on structure predictions it contains a 5’ stem-loop (containing the *sgrT* open reading frame), a 3’ stem-loop and an internal seed region (Figure 5A). To make use of the reusable part library, we generated two new level 0 parts, the first containing the PL promoter fused to the 5’ SgrS sequence (pSL133) and the second containing the 3’ SgrS sequence (pSL132). This strategy allowed the reuse of the SRL-library, which has been created for RybB (Figure 5B). The library of SgrS was created in parallel to the RybB library and treated in the same manner. In total, 380 candidates were screened for oxacillin susceptibility (Figure 5C) and subsequently sequenced in the same Nanopore sequencing run. Based on the results a defined SRL-library was obtained lacking the SRL of 0, 2, 26, 50 and 80 nt. The library was assessed on solid (data not shown) and in liquid media for oxacillin susceptibility. In contrast to the RybB library, there were only two lengths (36 and 42 nt) that showed a superior regulation at 50 µg/mL oxacillin (Figure 5D). In particular, SRL42 seems to have an increased repression of *acrA*, even in comparison to the tested RybB candidates. Notably, in the initial library we observed a sequence duplication in the downstream region of SgrS. However, it was experimentally proven that the sequence amplification did not have an influence (data not shown). Nevertheless, all subsequent experiments were performed with the corrected sequence for individually created synthetic SgrS sRNAs. We first tested the minimum SRL needed for functionality of synthetic SgrS sRNAs. As observed for RybB, we determined a minimal SRL of 12 nt for regulation of *acrA* using the oxacillin susceptibility assay on solid media (100 µg/mL oxacillin) (Figure 5E), which was also confirmed in liquid media at 75 µg/mL oxacillin (Figure S6).

**Figure 5.**
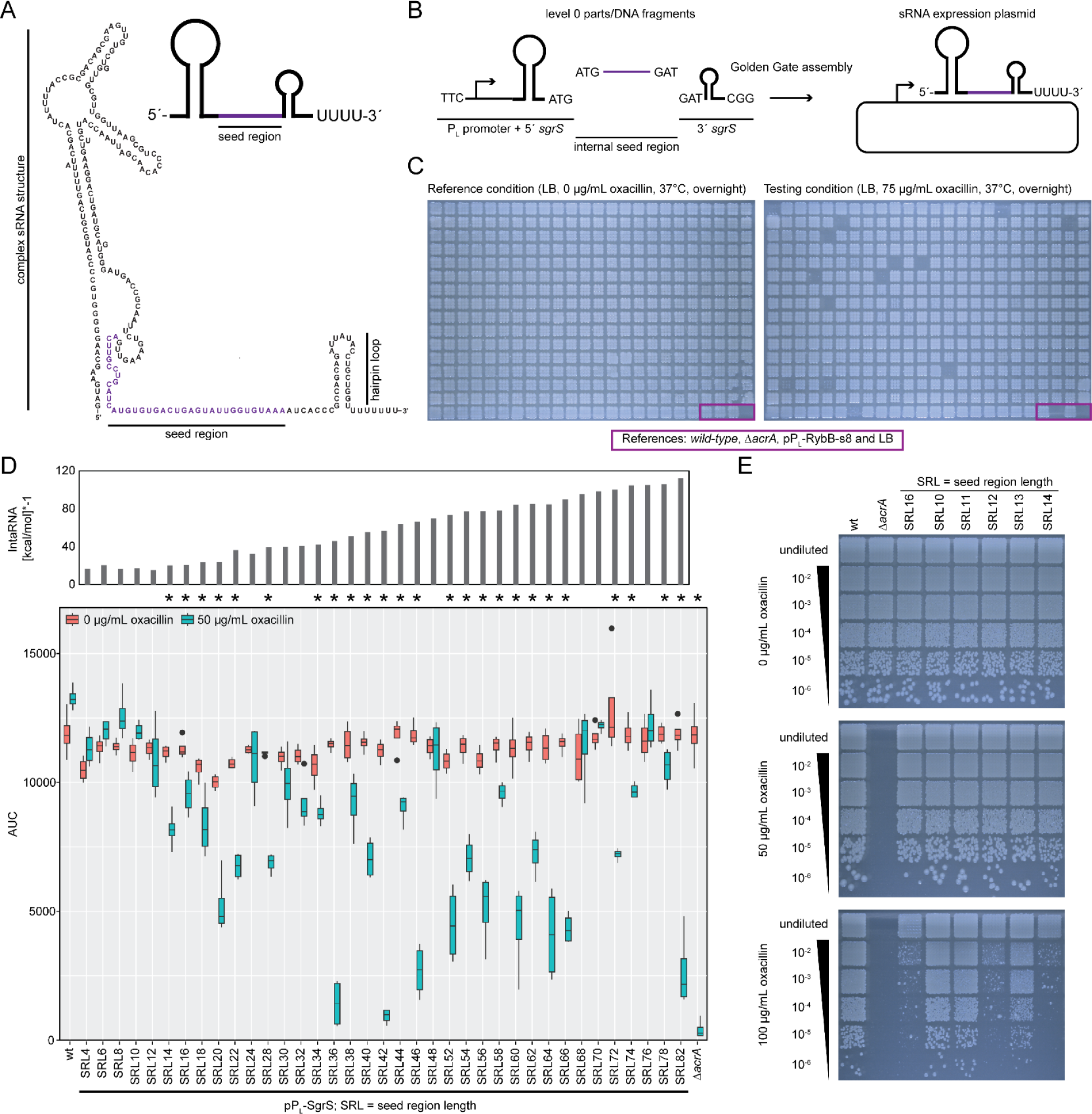
Adaptation of the SRL-library approach towards the SgrS sRNA for *acrA* targeting. **(A)** Predicted structure of SgrS according to RNAcentral (The RNAcentral Consortium, 2019) visualizes the complex 5’ secondary structure and the 3’ terminator stem-loop. For simplification a schematic is introduced for the subsequent figures. The seed region is indicated in purple and was adapted from Supplementary Figure S2 of Na *et al*. 2013 (Na et al., 2013). **(B)** Schematic of the Golden Gate assembly of the SRL-library using SgrS as scaffold. In contrast to RybB, the promoter sequence and the 5’ sequence of SgrS are fused allowing reuse of the PCR-amplified SRL-library. All other steps are identical to the cloning of RybB (Figure 1B). **(C)** Screening of the SgrS candidate library. 380 candidates and four respective controls were spotted from fresh overnight cultures into 3 × 3 grits on LB-agar plates with and without oxacillin. In contrast to the RybB candidate library, fewer candidates show reduced growth under the screening condition. Violet box indicates the position of indicated controls. **(D)** Liquid growth analysis indicates regulation of *acrA* for some of the selected candidates by accessing the area under the curve (AUC) in the presence and absence of oxacillin. According to the assay at least 14 nt are needed for regulation of *acrA*. Synthetic SgrS sRNAs with a SRL of 36 and 42 nt are the best performing. The SRL does not correlate with its functionality despite possessing lower binding energies. Binding energies are calculated by IntaRNA (for display reasons the values are multiplied by -1). Wild type (wt) and Δ*acrA* with an empty plasmid are serving as respective positive and negative controls. Oxacillin susceptibility assay was performed in quadruplicate. Student’s *t*-test was applied for statistical testing of the difference between growth with and without oxacillin (*: *P* < 0.05). **(E)** The minimum SRL for functionality of synthetic SgrS sRNAs was determined by oxacillin susceptibility assays on solid media. Similar to RybB, a SRL of 12 nt seems to be sufficient for regulation. A representative replicate is shown.

We were wondering why in particular SgrS with an SRL of 36 and 42 nt were performing superior in contrast to all other tested SgrS sRNAs. We used RNAfold (Lorenz et al., 2011) to predict the sRNA structures and observed that in most cases the seed region is sequestered (data not shown). In contrast, the seed regions of SRL36 and SRL42 are less likely engaged in intramolecular base-pairing. We further predicted the structures for SgrS sRNAs that deviate by 1 nt from SRL36 and SRL42. According to these predictions, the seed regions of SRL35 and SRL43 are likely accessible, while the seed regions of SRL37 and SRL41 are potentially occluded *via* intramolecular base-pairing (Figure S7). We hypothesized that SRL37 and SRL41 show reduced functionality due to the lack of seed region accessibility. To prove our hypothesis we constructed the respective synthetic SgrS sRNAs and tested their regulatory effect. Indeed the two candidates, which were predicted to contain a more accessible seed region, performed similarly to SRL36 and SRL42. In contrast, the regulatory potential of SRL37 and SRL41 was abolished or at least significantly impaired on solid media (Figure 6A). Interestingly, the differences in liquid growth conditions are less drastic, but are in line with the functionality of SRL37 on solid media at higher oxacillin concentrations (Figure 6B). Our results show that the accessibility of the sRNA seed region is an important design feature for the construction of functional sRNAs and that a difference of 1 nt may decide whether or not the synthetic sRNA is functional.

**Figure 6.**
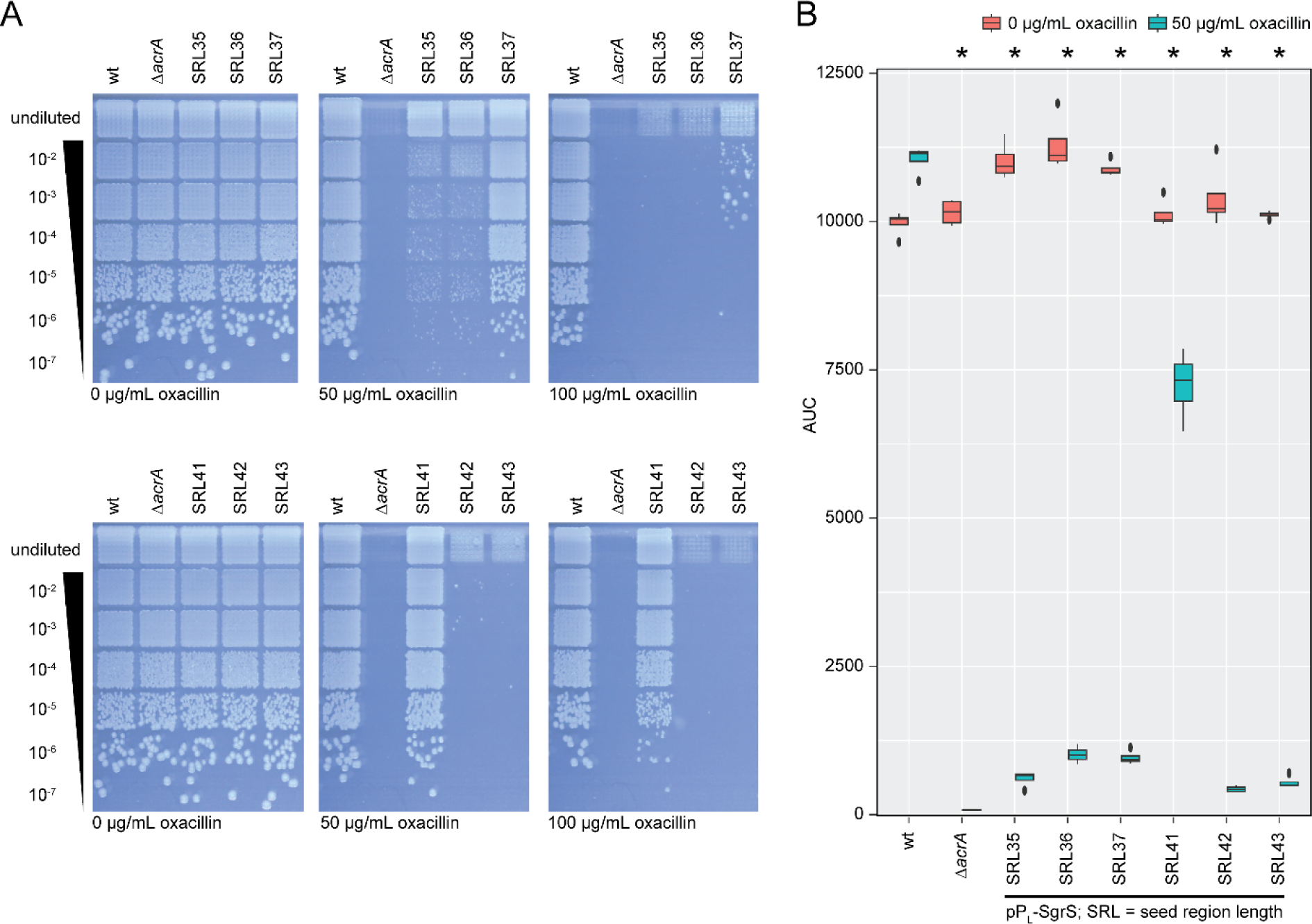
Synthetic SgrS sRNA functionality is affected by 1-nt changes. **(A)** Solid growth oxacillin susceptibility assay for SgrS SRL36 (upper panels) and SRL42 (lower panels) in comparison to synthetic SgrS sRNAs targeting *acrA* mRNA that deviate by 1 nt in their SRL. A representative replicate is shown. **(B)** Susceptibility test for the same strains as in (A) under liquid growth conditions. Oxacillin susceptibility assay was performed in quadruplicate. Student’s *t*-test was applied for statistical testing of the difference between growth with and without oxacillin (*: *P* < 0.05).

### An increasing seed region length helps to alleviate the Hfq-dependency of SgrS

The structurally complex sRNA SgrS was reported to depend on Hfq for riboregulation (Wadler and Vanderpool, 2007). When performing oxacillin susceptibility assays with the SgrS SRL-library in an *hfq* deletion background, we did not observe oxacillin-sensitive phenotypes on solid media (data not shown), suggesting that the synthetic SgrS sRNAs depend on Hfq for regulation of *acrA*. In liquid media however, increased sensitivity to oxacillin was observed for some SgrS variants in the Δ*hfq* strain, especially for those with an SRL above ∼50 nt (Figure S8). To further elucidate the impact of an increased SRL on the Hfq-dependency, we applied our fluorescent reporter assay (translational fusion of *acrA* with *syfp2*) for selected synthetic SgrS sRNAs. In the wild-type background, repression of the *acrA* reporter correlated well with the observed oxacillin sensitivity (*cf.* Figures 5D): variant SRL4 did not repress the reporter fusion, and variants SRL36 and SRL42 showed superior repression in comparison to SRL12, SRL66 and SRL82 (Figure 7A). In the Δ*hfq* background, the pattern changed, and enhanced repression was observed with an increasing SRL. Intriguingly, repression by SRL82 was comparable in both strain backgrounds (Figure 7A). In contrast to RybB, steady-state levels of the synthetic SgrS sRNAs were generally not reduced in the absence of Hfq, and some variants even showed increased levels (Figure 7B). Interestingly, the distinct processing pattern, which was observed for the RybB SRL-library, did not occur for SgrS. Our results indicate that an increasing seed region length helps to alleviate the Hfq-dependency of SgrS and that the position of the seed region (5’ end versus internal) impacts the SRL-dependent processing by RNase E.

**Figure 7.**
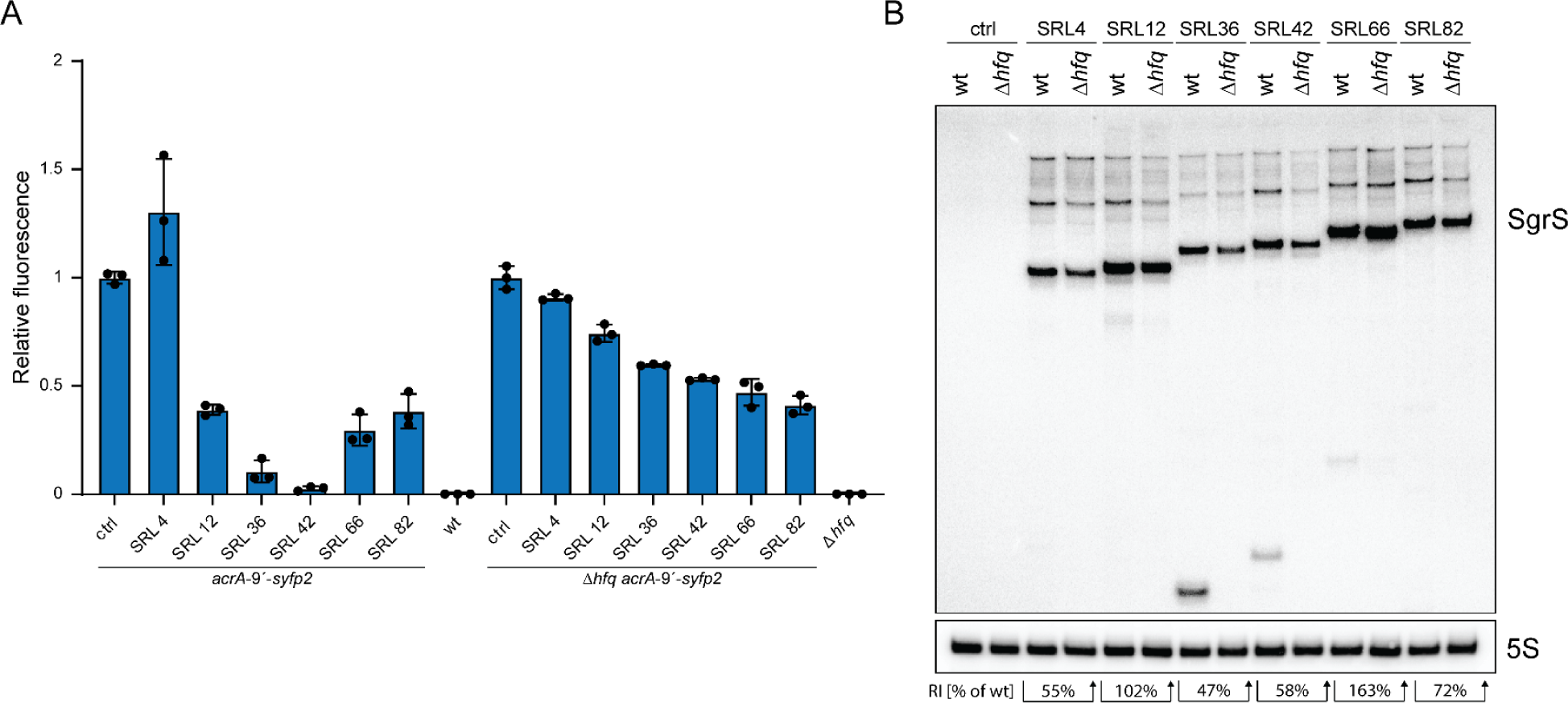
Regulation of *acrA* by selected synthetic SgrS sRNAs in wild type and Δ*hfq*. **(A)** Fluorescence of reporter strain *acrA-9’-syfp2* is normalized to the control strain, containing empty plasmid pSL009. In the wild type, reduction of fluorescence indicates sRNA functionality and correlates with phenotypes (*cf.* Figure 5D). In the absence of Hfq, the fluorescence intensity is generally increased. However, the Hfq dependence decreases with increasing SRL, and regulation by SgrS SRL82 is almost comparable between wild type and Δ*hfq*. The control (ctrl) refers to empty plasmid pSL009. A wild type (wt) and Δhfq strain without reporter construct were used as background controls, respectively. Fluorescence reporter assay was performed in triplicate. Student’s *t*-test was applied for statistical testing (*: *P* < 0.05). **(B)** The control (ctrl) refers to empty plasmid pSL009. 5S rRNA serves as a loading control. Abundance of SgrS sRNAs is only slightly reduced on average to 83% +/- 44% (RI = relative intensity in Δ*hfq* in comparison to wild type).

## Discussion

We established a workflow to benchmark and characterize synthetic sRNAs in high-throughput supported by laboratory automation equipment. We applied our previously developed assay for targeting of *acrA* mRNA, rendering *E. coli* cells susceptible towards the β-lactam antibiotic oxacillin (Köbel et al., 2022), in a library-based approach, and focused on the impact of the SRL on sRNA functionality. We combined the phenotypic assay with our recently described rapid and low-cost in-house long-read sequencing-based analysis workflow to connect phenotype and genotype (Ramirez Rojas et al., 2024a). Our presented library approach can rapidly identify synthetic sRNAs with improved functionality. Construction, testing and identification of optimal regulating sRNAs can be achieved within four to five days using the presented workflow (excluding sequence determination). To understand the mechanistic functions of synthetic sRNAs, this study went beyond the identification of improved synthetic sRNAs and characterized the library to improve our knowledge for future prediction, generation and application of synthetic sRNAs. A logical future iteration of the system could also allow the cloning of a UTR-library in front of the *acrA* reporter. This would facilitate the design and testing of synthetic sRNA regulators for virtually any gene of interest and sequence context. Furthermore, in combination with efficient barcode-based Nanopore sequencing, large seed- and UTR-libraries could be screened to generate sufficient training data for artificial intelligence design tools. Such a phenotype-based approach would complement fluorescence reporter assays and directly demonstrate sRNA functionality in a biological context.

Based on our results with RybB and SgrS, we conclude that a minimal SRL of 12 nt should be calculated for efficient regulation of target mRNAs. It may be important to avoid AU-rich sequences in short seed sequences to avoid RNase-mediated processing. A previous study has shown that an sRNA-mRNA interaction length of ≥ 15 nt was preferable when MicF was used as scaffold (Hoynes-O’Connor and Moon, 2016). However, efficient target repression was also achieved with some of the shorter MicF constructs (interaction length of < 15 nt). More importantly, the authors were able to show that the regulation strength saturated at an interaction length of ∼15 nt, and that synthetic sRNAs with an interaction length of ∼60 nt did not perform superior when compared to shorter sRNAs (Hoynes-O’Connor and Moon, 2016). This is in line with our findings that, in a wild-type background, increasing the SRL beyond the minimum of 12 nt does not necessarily improve regulatory performance of RybB. Since RybB and MicF are Hfq-dependent sRNAs, we conclude that synthetic seed regions as short as 12 nt are sufficient for efficient regulation as long as Hfq is present. Interestingly, it was observed for antisense PNAs conjugated to cell-penetrating peptides that 9-mer to 10-mer PNAs are sufficient for repression of target mRNAs (Goltermann et al., 2019; Popella et al., 2022). The slightly shorter minimal length that is needed for PNA-mediated repression might be founded in their higher intracellular stability and enhanced RNA-binding affinity (Pellestor and Paulasova, 2004). In the case of PNA-peptide conjugates, the PNA length needs to be minimized because it represents a constraint for the uptake into bacterial cells (Goltermann et al., 2019). In contrast, the SRL is not a limiting factor for endogenously produced synthetic sRNAs, suspending the need to design the shortest possible seed region.

We recently established a computational tool, SEEDling, for the selection of optimal seed regions based on structural considerations and potential off-target effects (Philipp et al., 2023), but we were lacking information about the optimal SRL. In a first attempt, we applied the tool to RybB sRNAs with 16-nt long seed regions, which is the native SRL of RybB (Brück et al., 2024). The results of our current study confirm that an SRL of 16 nt is well suited for RybB and probably other Hfq-dependent sRNAs with a 5’ seed region, such as MicA and MicF (Hoynes-O’Connor and Moon, 2016; Köbel et al., 2022). However, for other sRNAs, longer seed regions might be preferable for efficient target regulation. When transferring our SRL-library concept to the more complex sRNA SgrS (Wadler and Vanderpool, 2007), only two sRNAs (SRL36 and SRL42) showed strong repression of *acrA*. Hence, an SRL of above 35 nt was needed to achieve a performance similar to synthetic RybB sRNAs. In *E. coli*, native SgrS regulates the *ptsG* mRNA by imperfect base-pairing that involves 23 nt of the 31-nt long SgrS seed region (Maki et al., 2010; Vanderpool and Gottesman, 2004). Even though the SgrS SRL may be minimized to 14 nt without losing *ptsG* regulation (Maki et al., 2010), we conclude that SgrS relies on longer seed regions for efficient target regulation than other Hfq-dependent sRNAs (Santiago-Frangos and Woodson, 2018).

The structure of sRNAs is an important aspect to consider in the design process (Köbel et al., 2022; Philipp et al., 2023), which was particularly evident for synthetic SgrS sRNAs. One-nt changes may strongly impede functionality, as observed for SgrS SRL37 and SRL41 in comparison to SRL36 and SRL42, respectively. Based on secondary structure predictions we assume that synthetic SgrS sRNAs are sensitive to sequence changes that increase the local base-pairing probability within the seed region. Hence, in the case of SgrS, the overall structure may be even more important than the actual seed region. We suggest that SgrS seed regions should be designed with the aid of the SEEDling tool to control for structural changes that might affect seed region accessibility and, hence, impede regulation (Philipp et al., 2023).

Our data indicate that synthetic sRNAs with a 5’ seed region, such as RybB, are more likely prone to processing by the endoribonuclease RNase E, and that the synthetic seed region sequence may introduce AU-rich motives serving as cleavage sites. RNase E is known to target single-stranded RNAs at AU-rich sites in different bacteria (Chao et al., 2017; Forstner et al., 2018; McDowall et al., 1994). Our SRL-library fortuitously provides an increasing number of AU-rich motives with increasing SRL, generating up to four stable processing products for synthetic RybB sRNAs with an SRL ≥ 72 nt. However, our data does not indicate a decrease of RybB functionality with the appearance of additional processing sites. On the contrary, we conclude that processing of synthetic RybB sRNAs indicates successful target regulation due to coupled degradation of the sRNA-mRNA hybrid by RNase E (Prevost et al., 2011). It remains an open question whether the 5’ triphosphate of the RybB primary transcript needs to be converted into a 5’ monophosphate before processing can occur (Bandyra et al., 2012; Schilder and Gorke, 2023). The absence of distinct SgrS processing patterns further suggests that the synthetic SgrS seed regions are less likely to be accessible to RNase E as single-stranded regions, as also supported by our secondary structure predictions. Interestingly, the best performing SgrS sRNAs (SRL36 and SRL42) show at least one stable processing product, while the primary transcript is less abundant in the absence of Hfq. We conclude that these two sRNAs have a more single-stranded seed region, which allows efficient target regulation, but at the same time decreases stability.

With this study, we have extended the standardized Golden Gate basic-part toolbox for cloning of synthetic sRNAs. Golden Gate cloning has the advantage of rapid and reliable DNA assembly of the basic-part collection and is automation compatible (Engler et al., 2008; Köbel and Schindler, 2023; Weber et al., 2011). We have standardized our system using SapI as the Type IIS restriction enzyme, which creates 3-nt fusion sites, reducing the size of scars in the process of generating synthetic sRNAs, which is downstream fully compatible with the Modular Cloning standard of the group of Sylvestre Marillonnet (de Vries et al., 2024; Taylor et al., 2019). The development of a large standardized parts library for the generation of synthetic sRNAs can be a useful resource for the community in the future, allowing the rapid construction and characterization of synthetic sRNAs for basic and applied research. Larger datasets and the application of various sRNA scaffolds could generate an increased number of data points which may be used for machine learning approaches to predict the optimal synthetic sRNAs for their application in synthetic and engineering biology.

## Material and methods

### Strains and growth conditions used in this study

A full list of *E. coli* K-12 wild-type MG1655 and its derivatives used may be found in Table S1. *E. coli* K-12 wild-type MG1655 were used for physiological experiments. If not stated otherwise, strains were cultivated in LB medium in Erlenmeyer flasks at 37°C under continuous shaking at 180 rpm. If appropriate, antibiotics were present in the following concentrations: 100 μg/mL ampicillin (Amp), 50 μg/mL kanamycin (Kan), 25 μg/mL chloramphenicol (Cm), 120 µg/mL spectinomycin (Spec), and 10 μg/mL tetracycline (Tet).

### Oligodeoxynucleotides

All oligodeoxynucleotides were ordered from Integrated DNA Technologies (IDT, Coralville, USA) or Microsynth Seqlab (Göttingen, Germany) with standard desalting purification (Table S2). The oligonucleotide pool for the SRL library was ordered from Integrated DNA Technologies (IDT, Coralville, USA) in the form of an oPools™ oligo pool containing 42 distinct oligonucleotides (SLop2.1 to SLop2.42) at 50 pmol scale (Table S3). The primer used for the barcoding procedure for Nanopore sequencing were ordered in 96-well plates from Integrated DNA Technologies (IDT, Coralville, USA) with standard desalting purification (Ramirez Rojas et al., 2024a).

### PCR purification and plasmid extraction

If not indicated otherwise PCR amplicons and plasmid extractions were performed according to the open source magnetic bead procedure published by Oberacker *et al*. (2019) using SeraMag Speed Beads (Cytiva, Marlborough, USA) (Oberacker et al., 2019). For plasmid extractions cultures were grown in SMM medium (16 g/L tryptone, 10 g/L yeast extract, 5 g/L glycerol, and 1x M9 salts) with the appropriate antibiotic in 2-3 mL volume in tubes or 1 mL in 96-well deep-well plates (Köbel et al., 2022). Deep-well plates were inoculated in a Multitron HT shaker (Infors AG, Bottmingen, Switzerland) with 800 rpm and 80% humidity at 37°C overnight.

### Strain construction

Chromosomal manipulations were performed by λ red recombineering as described previously (Köbel et al., 2022). Briefly, for gene deletions, the chloramphenicol acetyltransferase (*cat*) gene was amplified to replace the gene(s) of interest. For the transcriptional *acrA* reporter fusion, the complete *syfp2* open reading frame together with an SD sequence was amplified and inserted immediately behind the *acrA* stop codon. Chromosomal constructs were verified by diagnostic PCR. All relevant primer pairs for λ red recombineering are provided in Table S2. Constructs were transduced to other strain backgrounds using P1 phages according to standard procedures (Lennox, 1955). Transductants were selected with chloramphenicol and verified by diagnostic PCR as before. If needed, resistance cassettes were removed by FRT-mediated flipping using plasmid 709-FLPe (Gene Bridges GmbH, Heidelberg, Germany) according to the manufacturer’s instructions.

### Generation of plasmids

Plasmids were mostly generated by Golden Gate cloning (all sequences are provided as GenBank files in Supporting Data S1). The generation of basic Golden Gate parts and the library construction is described in the following sections. Golden Gate assembly reactions were prepared with equimolar amounts for each DNA part and acceptor vector (20 to 40 fmol). Reactions were carried out in a thermocycler in 10 to 20 µL total volume with 1 μL of SapI and 1 μL of T4 DNA ligase in 1x T4 DNA ligase buffer and incubated with the following program: 50 cycles of 4 min at 16°C, 3 min at 37°C followed by 10 min at 50°C and 10 min at 80°C, and storage at 4°C. 10 µL of Golden Gate reactions were transformed into RbCl competent cells according to the standard procedure and candidates were selected by overnight cultivation on appropriate LB agar plates. A complete list of plasmids is provided in Table S4.

### Level 0 part generation

Level 0 parts were constructed by blunt end ligation as described before (de Vries et al., 2024) and all resulting plasmid files are provided as GenBank files in Supporting Data S1. Briefly, the primer for the PCR amplicons contained the corresponding overhangs and either the oligonucleotides or the obtained amplicon was phosphorylated using the T4 polynucleotide kinase (PNK) (NEB). The level 0 acceptor plasmid was amplified with the primer pair SLo0765/SLo0766 using Q5 DNA Polymerase (NEB) followed by a DpnI (NEB) digest prior ligation. The phosphorylated amplicon and the PCR amplified vector were ligated using T4 ligase for 2 h at room temperature in a 4:1 ratio. Transformation was performed into in house generated RbCl competent *E. coli* Top10 cells (Hanahan, 1983; Köbel et al., 2022). Selection was performed overnight on LB spectinomycin plates at 37°C. Level 0 parts were identified by colony PCR, followed by plasmid extraction and subsequent validation by external Sanger sequencing services (Microsynth, Göttingen, Germany).

### Library generation

Double stranded DNA was generated from the oligonucleotide pool by PCR using the primer SLo1505/1506 and Q5 DNA Polymerase with the following settings: Initial denaturation 0:20 min 98°C, followed by 35 PCR cycles [0:20 min 98°C, 0:20 min 62°C, 0:10 min 72°C] and final storage at 8°C. One fmol of the oligonucleotide pool was used in the reaction as a template. The RybB and SgrS plasmid libraries were generated by standard Golden Gate assembly combining the purified amplicon-library with pSL135 and pSL123 (for RybB) or pSL133 and pSL132 (for SgrS) into pSL137. The Golden Gate reaction was transformed into in house generated RbCl competent *E. coli* K-12 MG1655 cells (Hanahan, 1983; Köbel et al., 2022). The cell suspension was plated in dilution steps onto multiple 120 × 120 mm square petri dishes and incubated until single colonies appeared. 380 single colonies were picked onto 384-well microtiter plates with LB containing kanamycin utilizing the PIXL colony picking robot (Singer Instruments, Somerset, UK). Microtiter plates were incubated overnight on an Multitron HT (Infors AG, Bottmingen, Switzerland) incubator at 37°C with 500 rpm with 3 mm shaking and 80% relative humidity, only covered with the lid. The plates were used to inoculate liquid cultures in 384 well format for cryo-preservation and initial screening after overnight incubation with a Rotor HDA+ screening robot (Singer Instruments, Somerset, UK).

### Phenotypic screening and analysis of generated libraries

The initial screening of the 380 candidate sized library and 4 controls was done using overnight cultures grown on a Multitron HT (Infors AG, Bottmingen, Switzerland) at 500 rpm with 3 mm shaking, 80% humidity at 37°C. A Rotor HDA+ screening robot was used to spot the library into a 3 × 3 grid using 384-pin pads onto LB kanamycin media with varying oxacillin concentrations (0 to 200 µg/mL). Data was documented with the PhenoBooth (Singer Instruments, Somerset, UK). The determined length library for RybB and SgrS of constitutively expressed synthetic sRNA constructs was performed based on manual dilution series of indicated overnight cultures generated in 96-well or 384-well flat bottom microtiter plates using multichannel pipettes. A Rotor HDA+ screening robot (Singer Instruments, Somerset, UK) was afterwards used to transfer droplets onto solid agar plates containing LB kanamycin, and the indicated oxacillin concentrations. Programs either transfer 49 droplets for each dilution into a 7 × 7 grid (96-well source plates) or 9 droplets for each dilution into a 3 × 3 grid (384-well source plates), using 96-pin pads and 384-pin pads, respectively. Plates were incubated at 37°C overnight, and images were taken with the PhenoBooth plate documentation system (Singer Instruments, Somerset, UK).

Phenotypic screening in liquid media was performed in CLARIOstar plate readers (BMG Labtech, Ortenberg, Germany) equipped with plate holders conceptualized for kinetic measurements with constant shaking. Briefly, *E. coli* precultures were inoculated in microtiter plates (polystyrene, flat bottom; Greiner Bio-One, Kremsmünster, Austria) overnight at 37°C, 1000 rpm with 75% relative humidity in 100 μL of LB medium containing kanamycin on an Multitron HT incubator (Infors AG, Bottmingen, Switzerland). Assay cultures were inoculated from the overnight cultures with the Rotor HDA+ screening robot (Singer Instruments, Somerset, UK) into 96-well microtiter plates containing 200 μL of the respective LB medium supplemented with kanamycin and indicated oxacillin concentrations (0 to 100 μg/mL). Plates were sealed with optical clear sealing film with a PlateLoc plate sealer (Agilent, Santa Clara, USA). Measurements were taken with the following program: 120 s orbital shaking (1 mm amplitude) and 120 s linear shaking (1 mm amplitude), followed by wavelength measurement at 600 nm for 24 h. The program was compared to our previously described method using TopSeal-A PLUS (PerkinElmer, Waltham, USA) and Infinite M Nano+ plate reader (Tecan, Männedorf, Switzerland) (Köbel et al., 2022), no differences were observed. Data was processed with the Growthcurver R package (Sprouffske and Wagner, 2016) and visualized with custom R scripts. AUCs were calculated based on data points from t = 0 to t = 8 h for maximum dynamic range to compare growth.

### Nanopore sequencing and analysis of generated libraries

The procedure was performed according to Ramírez-Rojas *et al*. 2024. Briefly, amplicons were directly generated from the obtained colonies by a two-step PCR procedure (for details see (Ramirez Rojas et al., 2024a; Ramirez Rojas et al., 2024b). The first step attached universal sequences which are used in the second step as binding sequence for the barcoding primers. The amplified, dual barcoded amplicons of the RybB and SgrS candidates were pooled and purified with two other, similar libraries resulting in 1536 barcoded samples. One microgram of the purified library was used as input for the ligation sequencing kit SQK-LSK109 (Oxford Nanopore Technologies). The procedure was carried out according to the manufacturer guidelines and the library was sequenced on two Flongle flow cells FLO-FLG001 (R9.4.1) generating a total output of 0.9 Gb. Basecalling of the raw data was performed using ONT Guppy basecalling software (version 6.0.1). The amplicons were demultiplexed using MiniBar (Krehenwinkel et al., 2019), subsequently mapped against the references with minimap2 (Li, 2018), analyzed using Samtools (Danecek et al., 2021) and Integrative Genomics Viewer (Robinson et al., 2011). The procedure was later developed in an automated analysis pipeline described in (Ramirez Rojas et al., 2024a; Ramirez Rojas et al., 2024b).

### Northern blot analysis

For Northern blot analysis of synthetic sRNAs, expression plasmids were transformed to the indicated strain backgrounds. Total RNA was extracted from exponential-phase cultures (OD600 of approx. 0.4) using the hot acid-phenol method as described (Berghoff et al., 2017). Northern blot analysis was performed according to a standard protocol (Edelmann and Berghoff, 2019). In short, 10% polyacrylamide gels containing 1x TBE and 7 M urea were used for analysis of sRNAs. After gel separation, RNA was transferred to nylon membranes by semi-dry electroblotting. Oligodeoxynucleotides (Table S2) for detection of specific RNA species were end-labeled with [γ- ^32^P]-ATP using T4 polynucleotide kinase (PNK) (NEB). Prehybridization and hybridization were performed in Church buffer [0.5 M phosphate buffer (pH 7.2), 1% (w/v) bovine serum albumin, 1 mM EDTA, and 7% (w/v) SDS] at 42°C. After washing of membranes, phosphorimaging was applied for visualization using a Molecular Imager FX (Bio-Rad, Hercules, CA, USA). Signal intensities were quantified using the Quantity One 1-D Analysis Software (Version 4.6.6) (Bio-Rad, Hercules, CA, USA). sRNA signal intensities were normalized to the corresponding 5S rRNA signals.

### sRNA binding affinity prediction

The interaction energy for the *acrA* mRNA and the sRNA variants was calculated with IntaRNA (Mann et al., 2017) version 3.2.0 using the Vienna RNA package 2.4.14 and the “Turner 2004” energy model. IntaRNA was used in the heuristic mode (H) with the seed-extension strategy (model X) with lonely base pairs and GUs at helix ends allowed.

### Measurement of sYFP2 fluorescence

Chromosomal *syfp2* reporter strains, containing sRNA expression plasmids, were used for sYFP2 measurements as described previously (Brück et al., 2024; Köbel et al., 2022). Briefly, stationary-phase cultures were diluted 100-fold into LB medium, and 150 μL was loaded into wells of a transparent 96-well plate (polystyrene, flat bottom; Greiner Bio-One, Kremsmünster, Austria). Cells were cultivated in an Infinite M Nano^+^ microplate reader (Tecan, Männedorf, Switzerland) at 37°C and orbital shaking with an amplitude of 3.5 mm. The optical density was measured at 600 nm (OD600), and the sYFP2 fluorescence was monitored using excitation and emission wavelengths of 510 and 540 nm, respectively. The gain was set to 100. Measurements from wells containing pure LB medium were used for background correction of OD600 and sYFP2 values. Finally, sYFP2 fluorescence values were normalized to the corresponding OD600. Three biological replicates were obtained for each strain.

### Quantification and statistical analysis

For AUC and YFP-reporter measurements, Student’s *t*-test (unpaired, two-tailed: YFP values; paired, two-tailed: AUC values) was applied.

## Supporting information

- Supporting Information S1 - Figures S1 to S8, Tables S1 to S4, and supporting references.
- Supporting Data S1 - GenBank sequences for all generated plasmids.

## Resource availability

This study did not generate new unique reagents. Plasmids and strains generated in this study are available from the lead contact on request. All data are available within the study. This study did not generate new unique code. Any additional information required to reanalyze the data reported in this study is available from the lead contact upon request.

## Acknowledgments

This work was funded by the Max Planck Society in the framework of the MaxGENESYS project (D.S.), the European Union (NextGenerationEU) via the European Regional Development Fund (ERDF) by the state Hesse within the project “*biotechnological production of reactive peptides from waste streams as lead structures for drug development*” (D.S.), and an Exploration Grant from the Boehringer Ingelheim Foundation (BAB). J.G. was supported by the DFG (GE 3159/1-1). We are grateful to all laboratory members for extensive discussions on synthetic sRNAs. M.B. acknowledges that he was awarded the iFZ Masters 2023 Prize of the Justus Liebig University Giessen for this study.

## Author Contributions

B.A.B. and D.S. conceived the study. M.B., T.S.K., S.D., A.A.R.R., B.A.B., and D.S. conducted the experiments. M.B., J.G., B.A.B., and D.S. evaluated and analyzed the data. B.A.B. and D.S. wrote the manuscript with support of M.B. and J.G.. All authors read and approved the final manuscript.

## Declaration of interests

The authors declare no competing financial interest.

## Notes

### Competing Interest Statement

The authors have declared no competing interest.

