## Supporting Information S1 for "A library-based approach allows systematic and rapid evaluation of seed region length and reveals design rules for synthetic bacterial small RNAs"

**This file contains:**

Supporting Figures S1 – S8

Supporting Tables S1 – S4

Supporting References

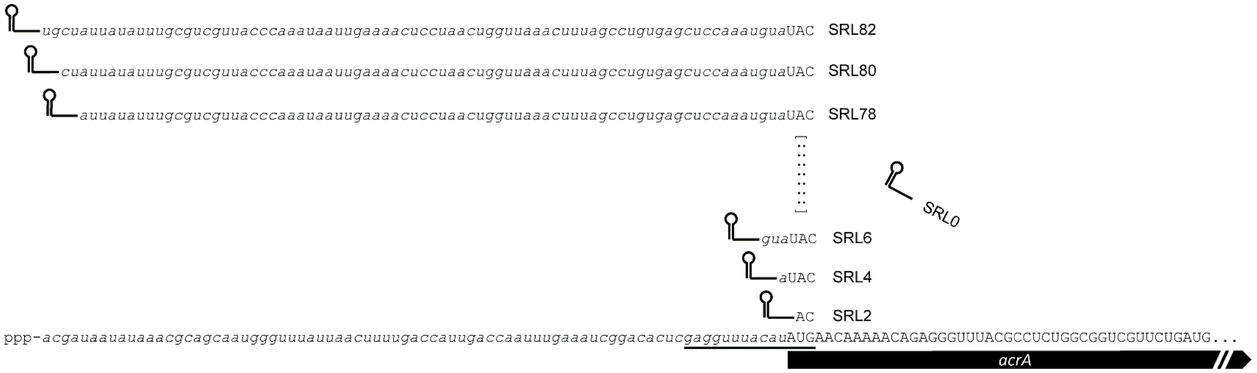


**Figure S1 | RybB seed region length library for targeting of *acrA* mRNA.** The *acrA* 5' UTR has a length of 79 nt, indicated by lowercase letters. The *acrA* open reading frame is indicated by the black arrow and capital letters. The RybB seed region length (SRL) library includes the start codon and extends to the 5' end of the transcript in 2-nt increments. The library consists of 42 different lengths, SRL0 represents a RybB scaffold that has no seed region and cannot bind to the *acrA* mRNA. The smaller sRNAs presumably do not bind, but are indicated at their cognate position to visualize the concept of the SRL library. For convenience, only SRL0 to SRL6 and SRL78 to SRL82 are visualized (dots indicate increasing seed region lengths). The translation initiation region (TIR) is underlined in the *acrA* mRNA. The *acrA* mRNA is truncated for visualization.

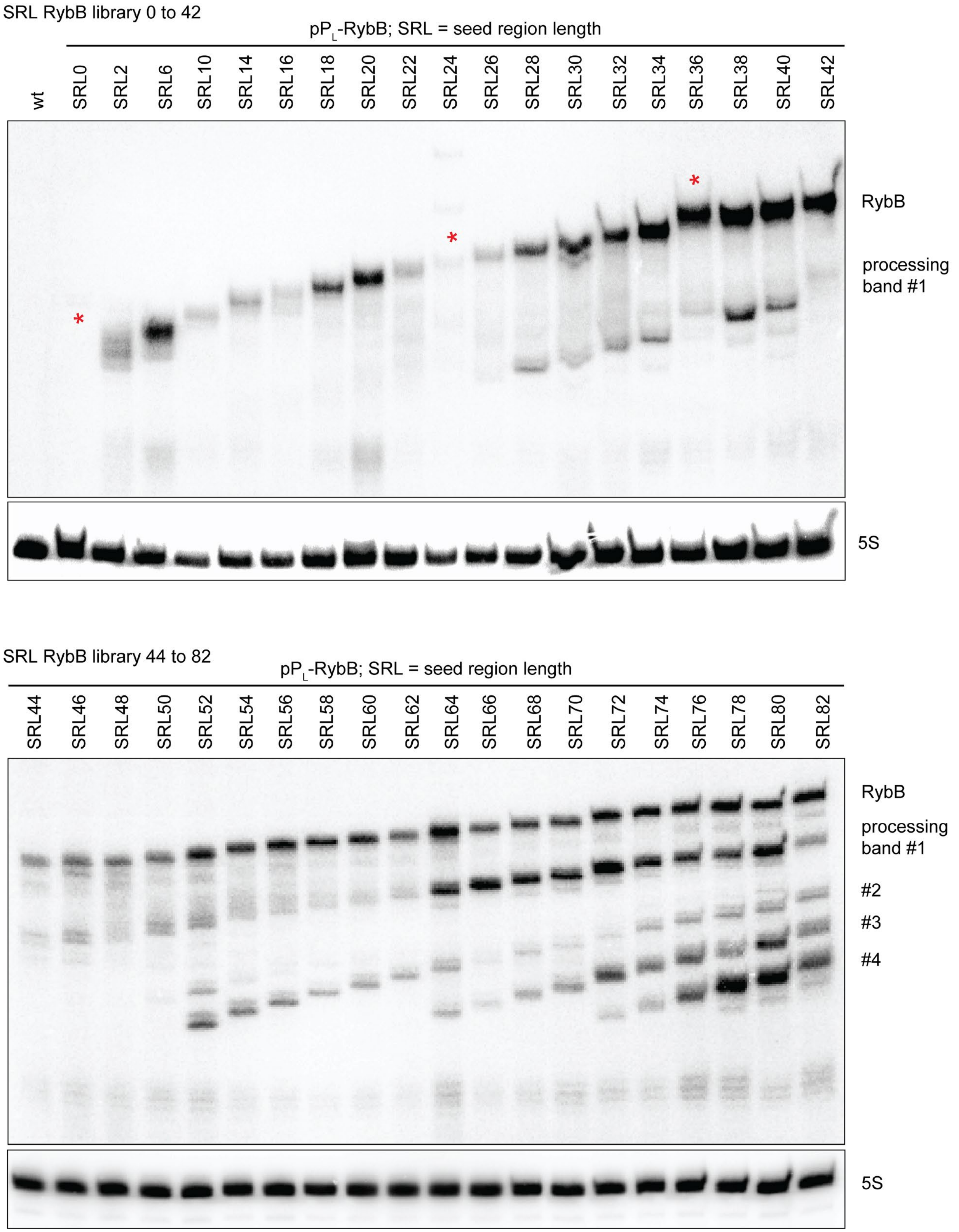


**Figure S2 | Northern blot analysis of the RybB SRL-library.** Northern blot analysis reveals a distinct processing pattern with the first band appearing with SRL26, the second with SRL52, the third with SRL64 and the fourth with

SRL72. Intensities of synthetic sRNAs and processing patterns do not correlate with functionality (*cf*. Figure 2B). Red asterisk indicate sRNAs which are not expressed (SRL0), are potentially a mixed population (SRL24) or are the wrong size (SRL36). 5S rRNA serves as a loading control.

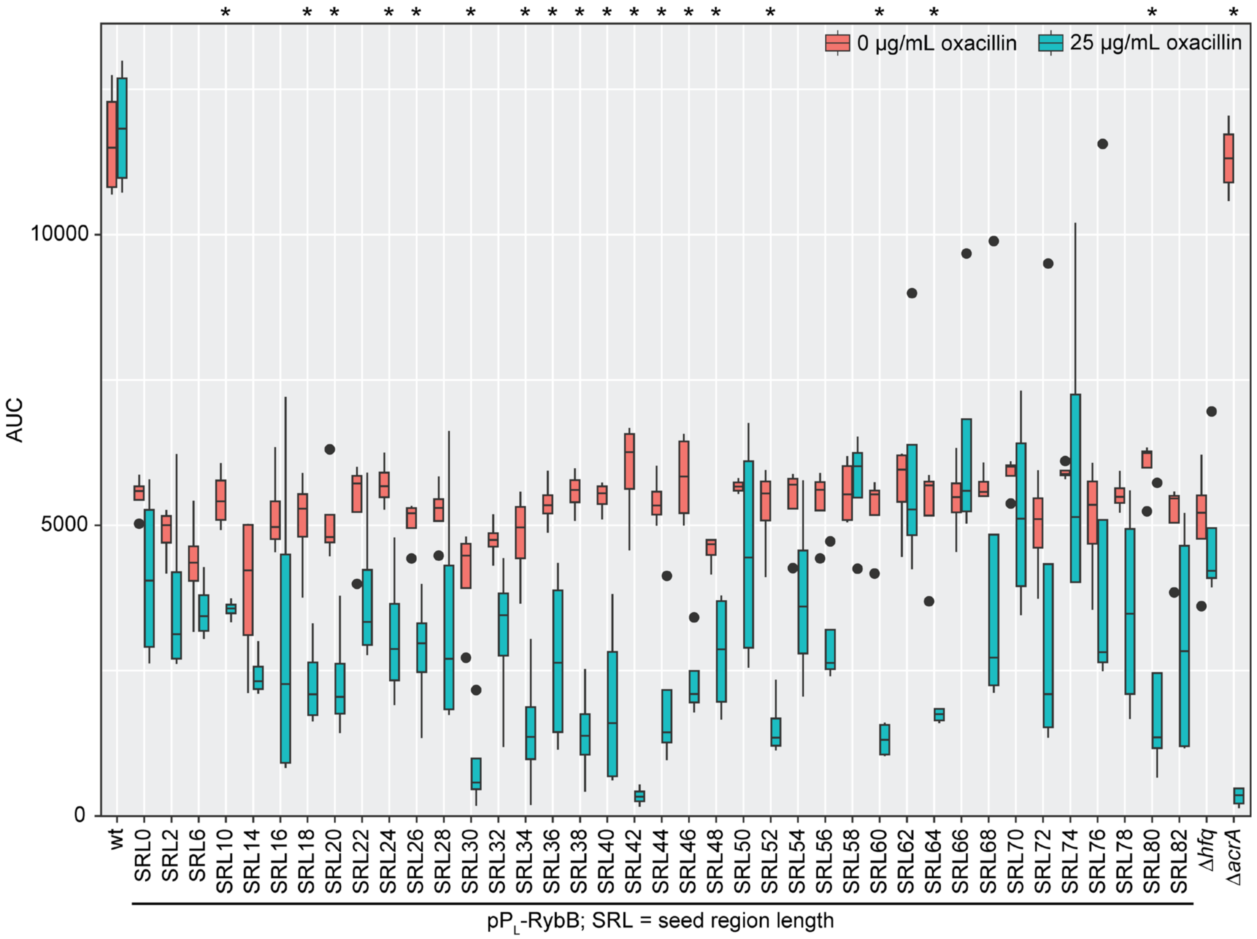


**Figure S3 | Determination of synthetic RybB sRNAs functionality in Δ*hfq*.** Liquid growth analysis of the SRL-library indicates no clear regulation for the synthetic sRNAs by accessing the area under the curve (AUC) in the absence and presence of oxacillin (25 µg/mL). The Δ*hfq* strain has a reduced viability, the AUC is decreased by approximately 2-fold even in the absence of oxacillin when compared to the wild type (wt). An increased SRL of the sRNAs does not show an effect. Wild type (wt), Δ*hfq* and Δ*acrA* with an empty plasmid serve as positive and negative controls. Oxacillin susceptibility assay was performed in quadruplicate. Student's *t*-test was applied for statistical testing against the Δ*hfq* reference strain (*: $P < 0.05$).

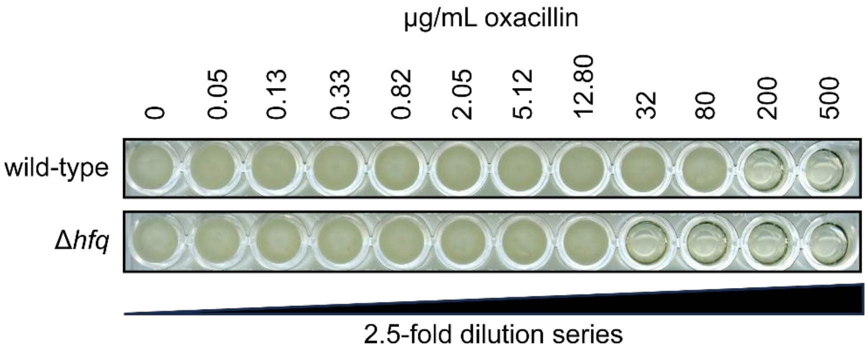


**Figure S4 | MIC determination for the *hfq* deletion strain.** Stationary-phase cultures were diluted 1,000-fold and loaded into 96-well plates. Oxacillin was present at the indicated concentrations (2.5-fold dilution series starting at 500 µg/mL). A well without oxacillin was used as growth control. The 96-well plates were incubated at 37°C under continuous shaking for 24 h. A representative experiment is shown.

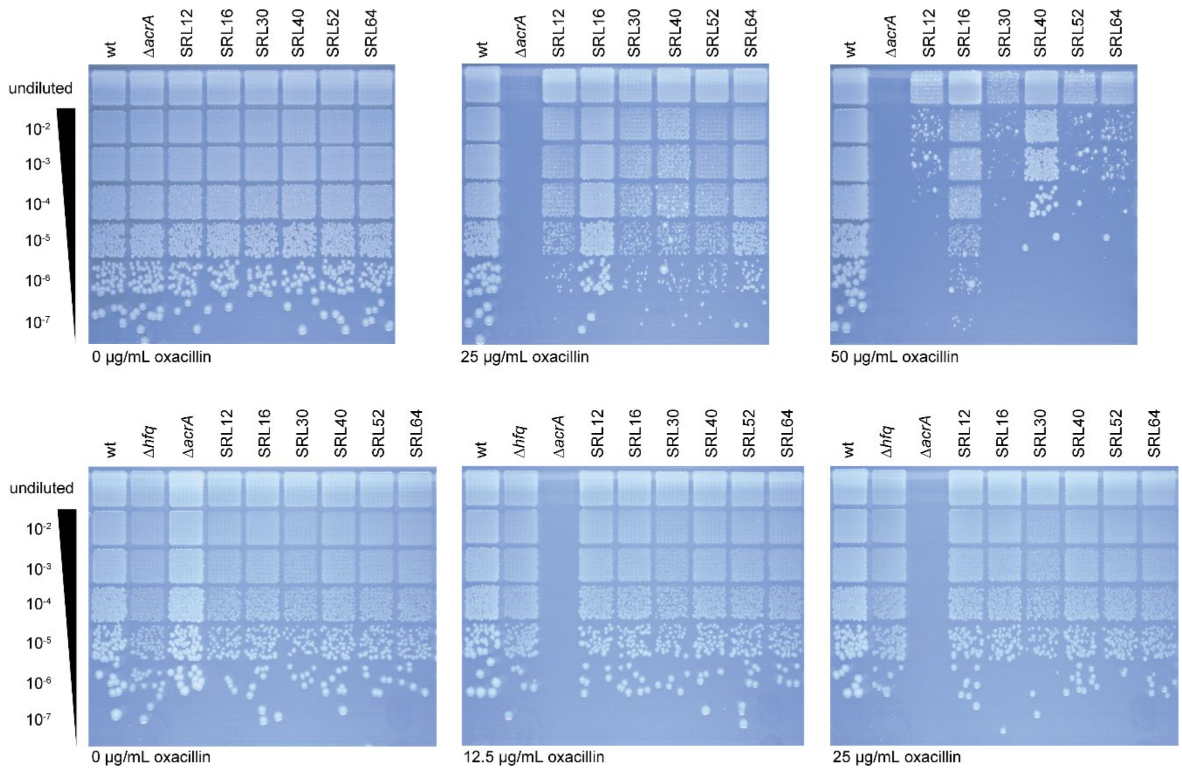


**Figure S5 | Solid growth oxacillin susceptibility assay for selected candidates in wild type and Δ*hfq*.** Upper panels show sRNA functionality in the wild type (wt). The lower panels show oxacillin susceptibility for Δ*hfq* expressing the indicated synthetic RybB sRNAs. No regulation of the *acrA* target can be observed in Δ*hfq*, which is consistent with the liquid media susceptibility assay (Figure 3A). Based on the reduced viability of Δ*hfq* in contrast to the wild type, the concentration of oxacillin for Δ*hfq* assays was reduced by half (Figure S4). A representative replicate is shown.

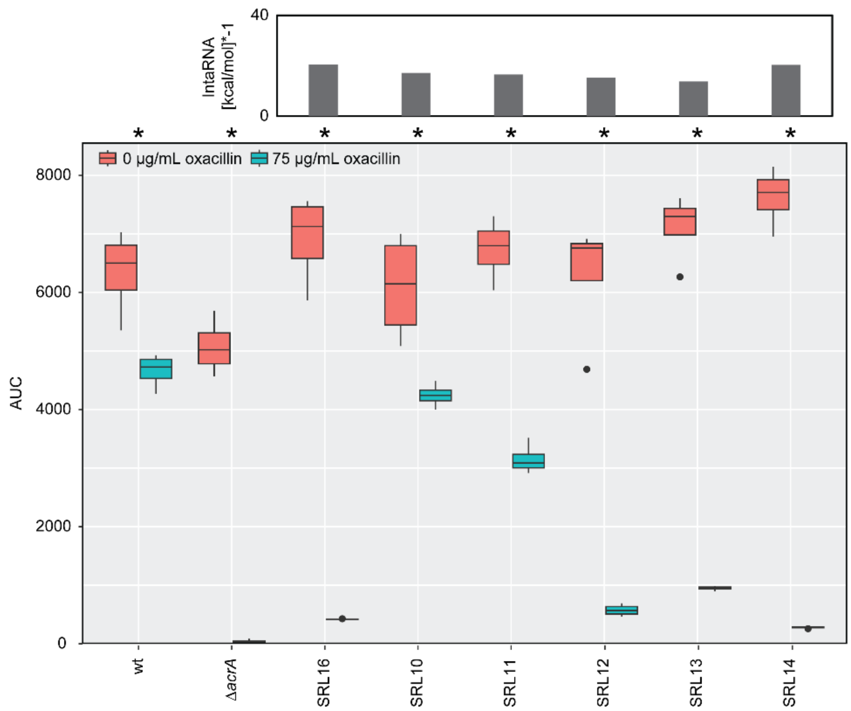


**Figure S6 | Liquid oxacillin susceptibility assay for minimal seed region length of synthetic SgrS sRNAs.** Oxacillin susceptibility assay was performed in quadruplicate. Student's *t*-test was applied for statistical testing of the difference between growth with and without oxacillin (*: $P < 0.05$).

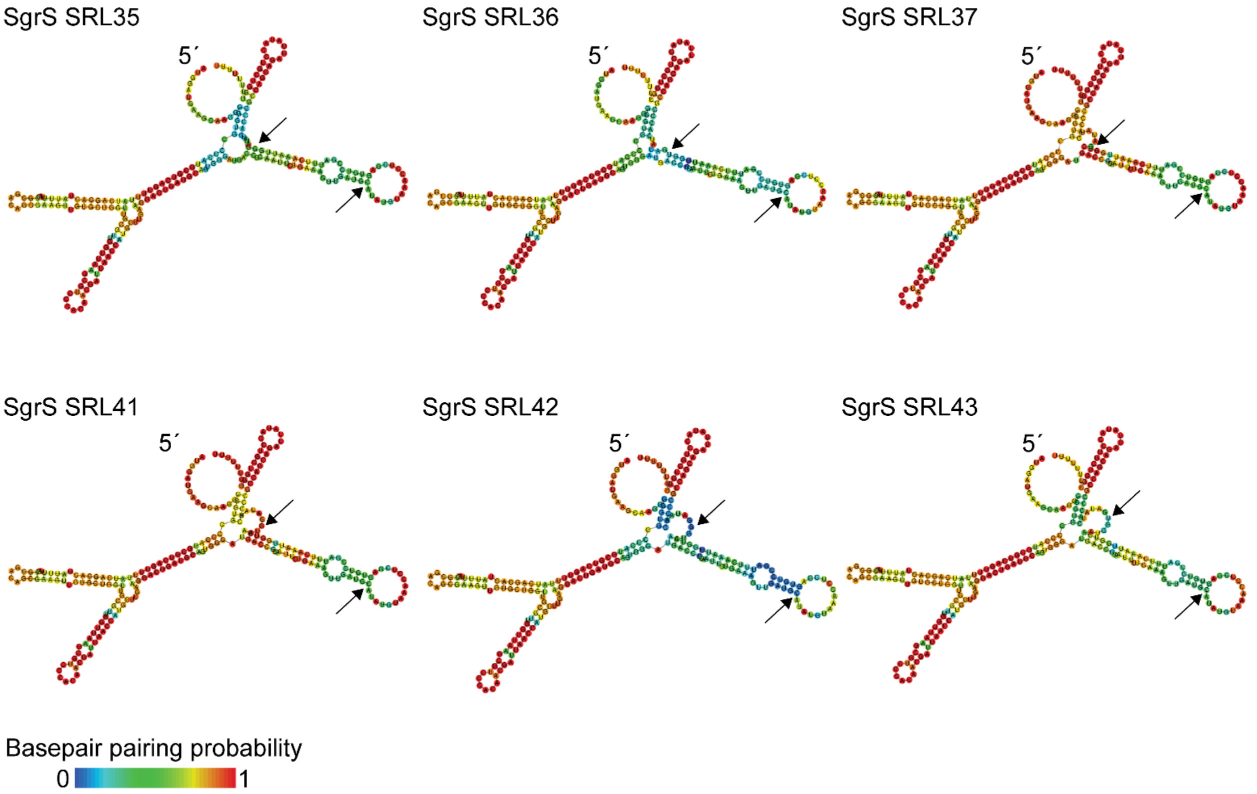


**Figure S7 | Structure predictions of selected synthetic SgrS sRNAs.** The seed regions of SgrS SRL35-37 and SRL41-43 are in a flexible stem loop structure with a comparably low base-pairing probability (seed regions are indicated by black arrows). The stem-loop structures of SgrS SRL37 and 41 have a higher base-pairing probability than the remaining SgrS sRNAs. Structures were generated using RNAfold (Lorenz et al., 2011).

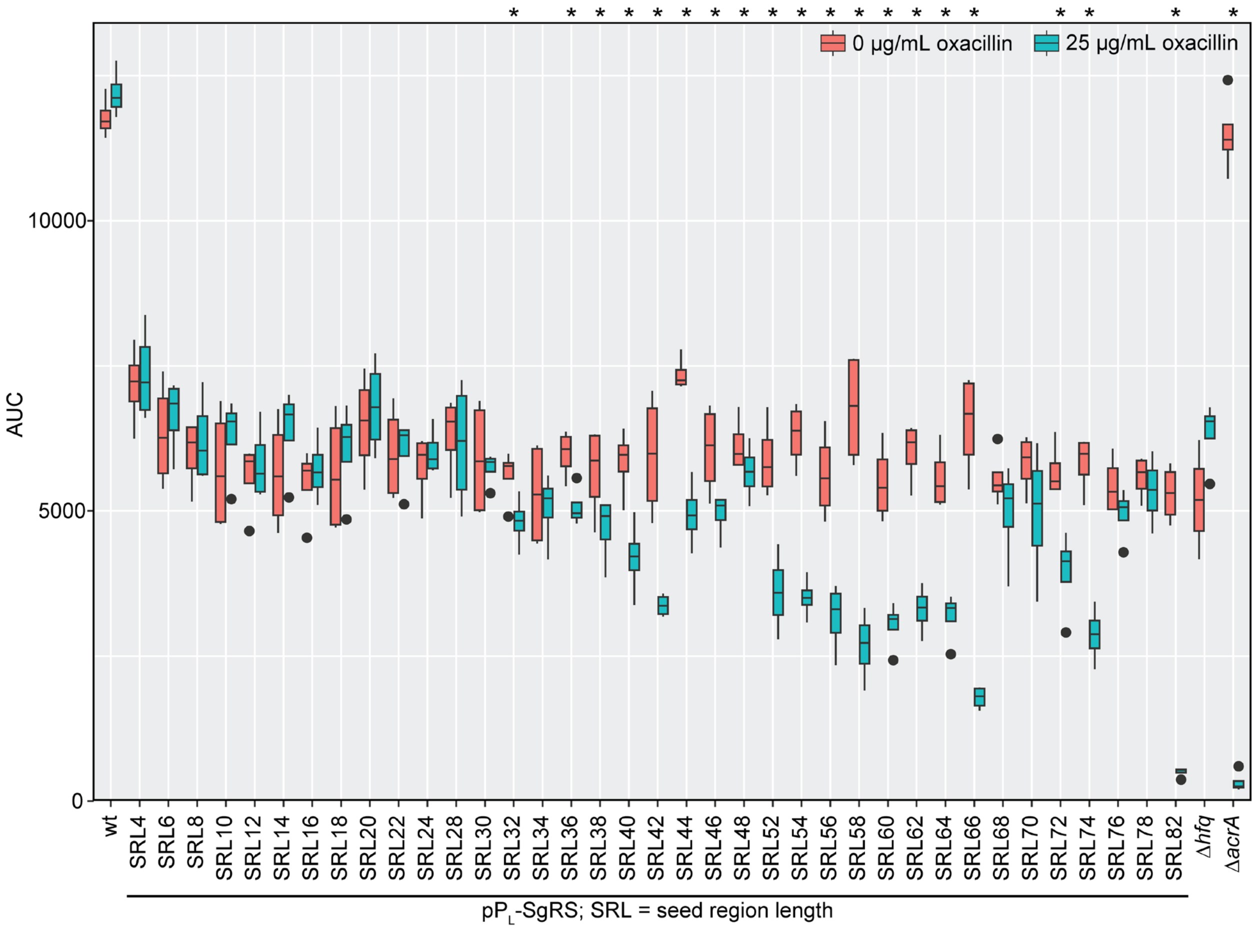


**Figure S8 | Determination of synthetic SgrS sRNAs functionality in Δ*hfq*.** Liquid growth analysis of the SgrS SRL-library indicates no strong regulation for the synthetic sRNAs by accessing the area under the curve (AUC) in the absence and presence of oxacillin (25 μg/mL). Increasing the SRL shows a mild correlation with reduced AUC in the presence of oxacillin. Notably, the best performing SgrS sRNAs (SRL36 and SRL42) are not the best regulating sRNAs in the absence of Hfq. Wild type (wt), Δ*hfq* and Δ*acrA* with an empty plasmid serve as respective positive and negative controls. Oxacillin susceptibility assay was performed in quadruplicate. Student's *t*-test was applied for statistical testing of the difference between growth with and without oxacillin (*: $P < 0.05$).

### Table S1 | Strains used in this study.

| Name | Relevant features | Reference |
|---|---|---|
| *E. coli* K-12 MG1655 | K-12 $F^-$ $\lambda^-$ | (Blattner et al., 1997) |
| *E. coli* DB3.1 | $F^-$ *gyrA462 endA1 glnV44* Δ(*sr1-recA*) *mcrB mrr hsdS20*($r_B^-$, $m_B^-$) *ara14 galK2 lacY1 proA2 rpsL20*($Str^R$) *xyl5* Δ*leu mtl1* | Invitrogen |
| *E. coli* Top10 | $F^-$ *mcrA* Δ(*mrr-hsdRMS-mcrBC*) φ80*lacZ*ΔM15 Δ*lacX74 nupG recA1 araD139* Δ(*ara-leu*)7697 *galE15 galK16 rpsL*($Str^R$) *endA1* $\lambda^-$ | Invitrogen |
| Δ*acrA* | MG1655 Δ*acrA*::*cat*, $Cm^R$ | (Köbel et al., 2022) |
| Δ*hfq* | MG1655 Δ*hfq*::*cat*, $Cm^R$ | this study |
| Δ*rybB* | MG1655 Δ*rybB*::*cat*, $Cm^R$ | (Köbel et al., 2022) |
| *acrA-9′-syfp2* | MG1655 *acrA-9′-syfp2-cat*, translational fusion of first 9 *acrA* codons to *syfp2*, $Cm^R$ | (Köbel et al., 2022) |
| Δ*hfq::FRT acrA-9'-syfp2-cat* | MG1655 Δ*hfq::FRT acrA-9'-syfp2-cat* | this study |
| *E. coli* BL321 | $RNaseIII^-$ (*rnc*$^-$) *nadB*$^+$ *purI*$^+$ | (Studier, 1975) |
| *E. coli* BL322 | $RNaseIII^+$ (*rnc*$^+$) *nadB*$^+$ *purI*$^+$ | (Studier, 1975) |
| *E. coli* N3431 | *lacZ43 relA1 spoT1 thi-1 rne-3071* | (Goldblum and Apririon, 1981) |
| *E. coli* N3433 | *lacZ43 relA1 spoT1 thi-1* | (Goldblum and Apririon, 1981) |
| N3431 Δ*rybB* | *lacZ43 relA1 spoT1 thi-1 rne-3071* Δ*rybB*::*cat*, $Cm^R$ | this study |
| N3433 Δ*rybB* | *lacZ43 relA1 spoT1 thi-1* Δ*rybB*::*cat*, $Cm^R$ | this study |

### Table S2 | Oligonucleotide sequences used and created in this study.

| ID | Sequence 5´ -> 3´ | Purpose |
|---|---|---|
| SLo0765 | TGAAGAGCAGGCACGAACCC | Forward primer to amplify level 0 plasmid for blunt cloning, based on e.g., pSL099; expected size 2.1 kb |
| SLo0766 | AGAAGAGCGAGCACAGAGTGC | Reverse primer to amplify level 0 plasmid for blunt cloning, based on e.g., pSL099; expected size 2.1 kb |
| SLo1503 | TTCTTGTGAGCGGATAACAATTGACATTGTGAGCGGATAACAAGATACTGAGCACCCATG | Forward primer for $P_L$ promoter generation in combination with SLo1504 for annealing into level 0 |
| SLo1504 | CATGGGTGCTCAGTATCTTGTTATCCGCTCACAATGTCAATTGTTATCCGCTCACAAGAA | Reverse primer for $P_L$ promoter generation in combination with SLo1503 for annealing into level 0 |
| SLo1505 | GGAAAGCTAGCTCTTCCATG | Forward primer to amplify oligonucleotide library with SapI site with primer SLo1506 |
| SLo1506 | CACAGCTTAGCTCTTCGATC | Reverse primer to amplify oligonucleotide library with SapI site with primer SLo1505 |
| SLo1510 | GATGTCCCCATTTTGTGGAG | Forward primer to amplify *rybB* scaffold and terminator with primer SLo1512 |
| SLo1511 | ATGGATGTCCCCATTTTGTGGAG | Forward primer to amplify *rybB* scaffold and terminator with primer SLo1512 |
| SLo1512 | CCGGGATGACGCTGCATTTTGTC | Reverse primer to amplify *rybB* scaffold and terminator with primer SLo1510 or SLo1511 |
| SLo1521 | ATGGATGAAGCAAGGGGGTGCCC | Forward primer to amplify wt *sgrS* and terminator with primer SLo1522 |
| SLo1522 | CCGTCATAAAAGCGACCAGCATAAATGC | Reverse primer to amplify *sgrS* scaffold and terminator with primer SLo1522, SLo1523 or SLo1524 |
| SLo1523 | GATATCACCCGCCAGCAGATTATACC | Forward primer to amplify *sgrS* scaffold and terminator with primer SLo1520 |
| SLo1524 | TCAACTTTCAGAATTGCGGTC | Overlap extension PCR primer with SLo1521 to generate subsequently with amplicon SLo1522/1525 for *sgrS* |
| SLo1525 | GACCGCAATTCTGAAAGTTGAATCACCCGCCAGCAGATTATACC | Overlap extension PCR primer with SLo1522 to generate subsequently with amplicon SLo1521/1524 for *sgrS* |
| SLo1526 | TTCTTGTGAGCGGATAACAATTGACATTGTGAGCGGATAACAAGATACTGAGCACCCATGGATGAAGCAAGGGGGTGCCC | Forward primer to generate promoter 5´ *sgrS* scaffold with SLo1527 |
| SLo1527 | CATCAACTTTCAGAATTGCGGTC | Reverse primer to generate promoter 5´-*sgrS* scaffold with SLo1526 |
| SLo1577 | CCCAGTCACGACGTTGTAAAACGCGTCAATTGTCTGATTCGTTACCA | Forward sRNA nanopore sequencing primer for barcoding; single TU constructs based on pSL137 – M13 barcoding compatible |
| SLo1578 | AGCGGATAACAATTTCACACAGGCTTCTCTCATCCGCCAAAACA | Reverse sRNA nanopore sequencing primer for barcoding; single TU constructs based on pSL137 – M13 barcoding compatible |
| SLo5132 | ATGCATATGTAAAC | Forward oligonucleotide for seed region with 11 nt |
| SLo5133 | ATCGTTTACATATG | Reverse oligonucleotide for seed region with 11 nt |
| SLo5134 | ATGCATATGTAAACC | Forward oligonucleotide for seed region with 12 nt |

| ID | Sequence 5´ -> 3´ | Purpose |
|---|---|---|
| SLo5135 | ATCGGTTTACATATG | Reverse oligonucleotide for seed region with 12 nt |
| SLo5136 | ATGCATATGTAAACCT | Forward oligonucleotide for seed region with 13 nt |
| SLo5137 | ATCAGGTTTACATATG | Reverse oligonucleotide for seed region with 13 nt |
| SLo5138 | ATGCATATGTAAACCTCGAGTGTCCGATTTCAAATTGG | Forward oligonucleotide for seed region with 35 nt |
| SLo5139 | ATCCCAATTTGAAATCGGACACTCGAGGTTTACATATG | Reverse oligonucleotide for seed region with 35 nt |
| SLo5140 | ATGCATATGTAAACCTCGAGTGTCCGATTTCAAATTGGTC | Forward oligonucleotide for seed region with 37 nt |
| SLo5141 | ATCGACCAATTTGAAATCGGACACTCGAGGTTTACATATG | Reverse oligonucleotide for seed region with 37 nt |
| SLo5142 | ATGCATATGTAAACCTCGAGTGTCCGATTTCAAATTGGTCAATG | Forward oligonucleotide for seed region with 41 nt |
| SLo5143 | ATCCATTGACCAATTTGAAATCGGACACTCGAGGTTTACATATG | Reverse oligonucleotide for seed region with 41 nt |
| SLo5144 | ATGCATATGTAAACCTCGAGTGTCCGATTTCAAATTGGTCAATGGT | Forward oligonucleotide for seed region with 43 nt |
| SLo5145 | ATCACCATTGACCAATTTGAAATCGGACACTCGAGGTTTACATATG | Reverse oligonucleotide for seed region with 43 nt |
| acrAB-scr-1 | GTATGTACCATAGCACGACG | Screening of *acrA* and *acrAB* manipulations |
| hfq-KO-1 | AAGGTTCAAAGTACAAATAAGCATATAAGGAAAAGAGAGATGTAGGCTGGAGCTGCTTC | Deletion of *hfq* by Lambda Red recombineering |
| hfq-KO-2 | AGGATCGCTGGCTCCCCGTGTAAAAAAACAGCCCGAAACCCTCATATGAATATCCTCCTTAGTTCC | Deletion of *hfq* by Lambda Red recombineering |
| hfq-scr-1 | GTTTATCGAGGTGTTTGCGC | Screening of *hfq* deletion |
| hfq-scr-2 | ATCACCTGCAATGCTTCGAC | Screening of *hfq* deletion |
| sYFP2_out | CGCGTCTTGTAGTTACCG | Screening of sYFP2 reporter gene |
| 5S probe-2 | CCTGGCAGTTCCCTACTCTCGCATGAGGAG | End-labeling for detection of 5S rRNA |
| RybB-probe-2 | GAAATGGCGGGGTTGATGGGCTCCACAAAATGGGGACATC | End-labeling for detection of RybB; binding after 16-nt seed region |
| SgrS-3'probe | AAAAAAAACCAGCAGGTATAATCTGCTGGCGGGTGAT | End-labeling for detection of SgrS (binds to 3' end) |

### Table S3 | Oligonucleotide sequences within the seed region oligo pool (SLop2).

| ID | Sequence 5´ -> 3´ * |
|---|---|
| SLop2.01 | ggaaagctaGCTCTTCc**ATGGAT**cGAAGAGCtaagctgtg |
| SLop2.02 | ggaaagctaGCTCTTCc**ATG**ca**GAT**cGAAGAGCtaagctgtg |
| SLop2.03 | ggaaagctaGCTCTTCc**ATG**cata**GAT**cGAAGAGCtaagctgtg |
| SLop2.04 | ggaaagctaGCTCTTCc**ATG**catatg**GAT**cGAAGAGCtaagctgtg |
| SLop2.05 | ggaaagctaGCTCTTCc**ATG**catatgta**GAT**cGAAGAGCtaagctgtg |
| SLop2.06 | ggaaagctaGCTCTTCc**ATG**catatgtaaa**GAT**cGAAGAGCtaagctgtg |
| SLop2.07 | ggaaagctaGCTCTTCc**ATG**catatgtaaacc**GAT**cGAAGAGCtaagctgtg |
| SLop2.08 | ggaaagctaGCTCTTCc**ATG**catatgtaaacctc**GAT**cGAAGAGCtaagctgtg |
| SLop2.09 | ggaaagctaGCTCTTCc**ATG**catatgtaaacctcga**GAT**cGAAGAGCtaagctgtg |
| SLop2.10 | ggaaagctaGCTCTTCc**ATG**catatgtaaacctcgagt**GAT**cGAAGAGCtaagctgtg |
| SLop2.11 | ggaaagctaGCTCTTCc**ATG**catatgtaaacctcgagtgt**GAT**cGAAGAGCtaagctgtg |
| SLop2.12 | ggaaagctaGCTCTTCc**ATG**catatgtaaacctcgagtgtcc**GAT**cGAAGAGCtaagctgtg |
| SLop2.13 | ggaaagctaGCTCTTCc**ATG**catatgtaaacctcgagtgtccga**GAT**cGAAGAGCtaagctgtg |
| SLop2.14 | ggaaagctaGCTCTTCc**ATG**catatgtaaacctcgagtgtccgatt**GAT**cGAAGAGCtaagctgtg |
| SLop2.15 | ggaaagctaGCTCTTCc**ATG**catatgtaaacctcgagtgtccgatttc**GAT**cGAAGAGCtaagctgtg |
| SLop2.16 | ggaaagctaGCTCTTCc**ATG**catatgtaaacctcgagtgtccgatttcaa**GAT**cGAAGAGCtaagctgtg |
| SLop2.17 | ggaaagctaGCTCTTCc**ATG**catatgtaaacctcgagtgtccgatttcaaat**GAT**cGAAGAGCtaagctgtg |
| SLop2.18 | ggaaagctaGCTCTTCc**ATG**catatgtaaacctcgagtgtccgatttcaaattg**GAT**cGAAGAGCtaagctgtg |
| SLop2.19 | ggaaagctaGCTCTTCc**ATG**catatgtaaacctcgagtgtccgatttcaaattggt**GAT**cGAAGAGCtaagctgtg |
| SLop2.20 | ggaaagctaGCTCTTCc**ATG**catatgtaaacctcgagtgtccgatttcaaattggtca**GAT**cGAAGAGCtaagctgtg |
| SLop2.21 | ggaaagctaGCTCTTCc**ATG**catatgtaaacctcgagtgtccgatttcaaattggtcaat**GAT**cGAAGAGCtaagctgtg |
| SLop2.22 | ggaaagctaGCTCTTCc**ATG**catatgtaaacctcgagtgtccgatttcaaattggtcaatgg**GAT**cGAAGAGCtaagctgtg |
| SLop2.23 | ggaaagctaGCTCTTCc**ATG**catatgtaaacctcgagtgtccgatttcaaattggtcaatggtc**GAT**cGAAGAGCtaagctgtg |
| SLop2.24 | ggaaagctaGCTCTTCc**ATG**catatgtaaacctcgagtgtccgatttcaaattggtcaatggtcaa**GAT**cGAAGAGCtaagctgtg |
| SLop2.25 | ggaaagctaGCTCTTCc**ATG**catatgtaaacctcgagtgtccgatttcaaattggtcaatggtcaaaa**GAT**cGAAGAGCtaagctgtg |
| SLop2.26 | ggaaagctaGCTCTTCc**ATG**catatgtaaacctcgagtgtccgatttcaaattggtcaatggtcaaaagt**GAT**cGAAGAGCtaagctgtg |
| SLop2.27 | ggaaagctaGCTCTTCc**ATG**catatgtaaacctcgagtgtccgatttcaaattggtcaatggtcaaaagtta**GAT**cGAAGAGCtaagctgtg |
| SLop2.28 | ggaaagctaGCTCTTCc**ATG**catatgtaaacctcgagtgtccgatttcaaattggtcaatggtcaaaagttaat**GAT**cGAAGAGCtaagctgtg |
| SLop2.29 | ggaaagctaGCTCTTCc**ATG**catatgtaaacctcgagtgtccgatttcaaattggtcaatggtcaaaagttaataa**GAT**cGAAGAGCtaagctgtg |

| ID | Sequence 5´ -> 3´ * |
|---|---|
| SLop2.30 | ggaaagctaGCTCTTCc**ATG**catatgtaaacctcgagtgtccgatttcaaattggtcaatggtcaaaagttaataaac**GAT**cGAAGAGCtaagctgtg |
| SLop2.31 | ggaaagctaGCTCTTCc**ATG**catatgtaaacctcgagtgtccgatttcaaattggtcaatggtcaaaagttaataaaccc**GAT**cGAAGAGCtaagctgtg |
| SLop2.32 | ggaaagctaGCTCTTCc**ATG**catatgtaaacctcgagtgtccgatttcaaattggtcaatggtcaaaagttaataaacccat**GAT**cGAAGAGCtaagctgtg |
| SLop2.33 | ggaaagctaGCTCTTCc**ATG**catatgtaaacctcgagtgtccgatttcaaattggtcaatggtcaaaagttaataaacccattg**GAT**cGAAGAGCtaagctgtg |
| SLop2.34 | ggaaagctaGCTCTTCc**ATG**catatgtaaacctcgagtgtccgatttcaaattggtcaatggtcaaaagttaataaacccattgct**GAT**cGAAGAGCtaagctgtg |
| SLop2.35 | ggaaagctaGCTCTTCc**ATG**catatgtaaacctcgagtgtccgatttcaaattggtcaatggtcaaaagttaataaacccattgctgc**GAT**cGAAGAGCtaagctgtg |
| SLop2.36 | ggaaagctaGCTCTTCc**ATG**catatgtaaacctcgagtgtccgatttcaaattggtcaatggtcaaaagttaataaacccattgctgcgt**GAT**cGAAGAGCtaagctgtg |
| SLop2.37 | ggaaagctaGCTCTTCc**ATG**catatgtaaacctcgagtgtccgatttcaaattggtcaatggtcaaaagttaataaacccattgctgcgttt**GAT**cGAAGAGCtaagctgtg |
| SLop2.38 | ggaaagctaGCTCTTCc**ATG**catatgtaaacctcgagtgtccgatttcaaattggtcaatggtcaaaagttaataaacccattgctgcgtttat**GAT**cGAAGAGCtaagctgtg |
| SLop2.39 | ggaaagctaGCTCTTCc**ATG**catatgtaaacctcgagtgtccgatttcaaattggtcaatggtcaaaagttaataaacccattgctgcgtttatat**GAT**cGAAGAGCtaagctgtg |
| SLop2.40 | ggaaagctaGCTCTTCc**ATG**catatgtaaacctcgagtgtccgatttcaaattggtcaatggtcaaaagttaataaacccattgctgcgtttatatta**GAT**cGAAGAGCtaagctgtg |
| SLop2.41 | ggaaagctaGCTCTTCc**ATG**catatgtaaacctcgagtgtccgatttcaaattggtcaatggtcaaaagttaataaacccattgctgcgtttatattatc**GAT**cGAAGAGCtaagctgtg |
| SLop2.42 | ggaaagctaGCTCTTCc**ATG**catatgtaaacctcgagtgtccgatttcaaattggtcaatggtcaaaagttaataaacccattgctgcgtttatattatcgt**GAT**cGAAGAGCtaagctgtg |

* underlined nucleotides = SapI recognition site, **bold nucleotides** = restriction site

**Table S4 | Plasmids used and created in this study, plasmid files except pSIM5 and 709-FLPe are available in GenBank format in Supporting Data S1.**

| ID | Relevant features | Parental plasmid | Reference |
|---|---|---|---|
| pSIM5 | λ red expression vector, pSC101 *ori*, *repA$^{ts}$*, Tet$^R$ | | (Datta et al., 2006) |
| 709-FLPe | FLPe expression plasmid, pSC101-ts *ori*, Amp$^R$ | | Gene Bridges GmbH |
| p-PL-RybB-s8 | RybB s8, Kan$^R$ | | (Köbel et al., 2022) |
| pSLcol_05 | RybB SRL library, Kan$^R$ | pSL137 | this study |
| pSLcol_08 | SgrS SRL library, Kan$^R$ | pSL137 | this study |
| pSL009 | Empty control vector, Kan$^R$ | pBAD | (Köbel et al., 2022) |
| pSL099 | Level 0 plasmid for subcloning of fragments to be released with SapI; Spec$^R$ | pMA60 (Schindler et al., 2016) | (Brück et al., 2024) |
| pSL123 | RybB scaffold with downstream region level 0 part; Spec$^R$ | pSL099 | this study |
| pSL132 | 3´ *sgrS* with downstream region level 0 part; Spec$^R$ (with downstream duplicated sequence) | pSL099 | this study |
| pSL133 | $P_L$ promoter + 5´ *sgrS* level 0 part; Spec$^R$ | pSL099 | this study |
| pSL135 | $P_L$ promoter level 0 part; Spec$^R$ | pSL099 | this study |
| pSL137 | Derivative of pBAD with SapI recognition site for Golden Gate cloning, Kan$^R$ | pBAD | (Köbel et al., 2022) |
| pSL571 | RybB SRL11, Kan$^R$ | pSL137 | this study |
| pSL572 | RybB SRL12, Kan$^R$ | pSL137 | this study |
| pSL573 | RybB SRL13, Kan$^R$ | pSL137 | this study |
| pSL598 | 3´ *sgrS* with downstream region level 0 part; Spec$^R$ (Duplicated downstream sequence removed) | pSL099 | this study |
| pSL723 | SgrS SRL11, Kan$^R$ | pSL137 | this study |
| pSL724 | SgrS SRL13, Kan$^R$ | pSL137 | this study |
| pSL725 | SgrS SRL35, Kan$^R$ | pSL137 | this study |
| pSL726 | SgrS SRL37, Kan$^R$ | pSL137 | this study |
| pSL727 | SgrS SRL41, Kan$^R$ | pSL137 | this study |
| pSL728 | SgrS SRL43, Kan$^R$ | pSL137 | this study |
